# Single-dose administration of therapeutic divalent siRNA targeting MECP2 prevents lethality for one year in an MECP2 duplication mouse model

**DOI:** 10.1101/2025.03.26.645328

**Authors:** Vignesh N. Hariharan, Ashley Summers, Amy E. Clipperton-Allen, Jillian Caiazzi, Samuel R. Hildebrand, Daniel O’ Reilly, Qi Tang, Zachary Kennedy, Dimas Echeverria, Nicholas McHugh, David Cooper, Jacqueline Souza, Chantal Ferguson, Laurent Bogdanik, Monica Coenraads, Anastasia Khvorova

## Abstract

MECP2 duplication syndrome (MDS) is a rare X-linked neurodevelopmental disorder caused by duplications of the dosage-sensitive methyl-CpG-binding protein 2 (MECP2) gene. Developing effective therapies for MDS is particularly challenging due to the variability in MECP2 expression among patients and the potential risk of inducing Rett syndrome through excessive pharmacological intervention. Reducing dosage to optimize silencing levels often compromises durability and necessitates increased dosing frequency. We present here a series of fully chemically modified small interfering RNAs (siRNAs) designed for both isoform-selective and total MECP2 silencing. Among these, we identify six lead siRNA candidates across two distinct chemical scaffolds, achieving targeted total MECP2 expression reductions ranging from 25% to 75%, sustained for at least four months following a single administration. The efficacy and safety of human ortholog silencing were evaluated using two mouse models with distinct levels of human MECP2 transgene expression. In the severe duplication model, a single dose of the total isoform-silencing siRNA fully rescued early mortality and behavioral impairments. Additionally, we show that the isoform-selective targeting strategy may be safer in mild cases of MDS where exaggerated pharmacology may lead to Rett Syndrome. Overall, this study introduces a series of preclinical candidates with the capacity to address the varying levels of MECP2 duplication encountered in clinical settings. Furthermore, it establishes a target selection strategy that may be applied to other dosage-sensitive gene imbalances.

**One Sentence Summary:** Therapeutic siRNAs provide safe and durable modulation of MECP2 for the treatment of mild and severe MECP2 Duplication Syndrome.

## INTRODUCTION

Methyl-CpG-binding protein 2 (MECP2) duplication syndrome (MDS) is a rare X-linked neurodevelopmental disorder resulting from the duplication of the Xq28 chromosomal region, which includes the dosage-sensitive MECP2 gene. MDS presents with a broad spectrum of severe clinical manifestations (*1*) with no cure for the disease. Mosaic loss-of-function mutations in MECP2 lead to Rett syndrome, highlighting the necessity of precise MECP2 dosage regulation for maintaining normal neurological function (*2*). Given the genetic basis of MDS, nucleic acid-based therapeutics (NATs), such as antisense oligonucleotides (ASOs) and small interfering RNAs (siRNAs), represent promising therapeutic avenues. These modalities enable precise modulation of gene expression by inducing transcriptional silencing through RNA interference (RNAi). Among these, an ASO named ION440 is the most advanced candidate currently under development for MDS, with a single dose providing up to twelve weeks of MECP2 modulation, with transcriptional correction for up to 16 weeks post administration. Clinical trials for this therapeutic candidate are anticipated to begin by the end of 2024 under the sponsorship of Ionis Pharmaceuticals (NCT06430385) (*3, 4*).

A key challenge in MECP2 therapeutics is precise dose titration to avoid excessive silencing, which may induce Rett syndrome (RTT). Female RTT patients, heterozygous for loss-of-function MECP2 mutations, exhibit a mosaic of wildtype and mutant *MECP2*-expressing cells (*5*), raising questions about whether uniform over-silencing using NATs could replicate RTT’s mosaic pathology. The pharmacodynamics of ASOs and siRNAs are inherently dose-dependent, with higher doses leading to greater tissue accumulation and sustained gene silencing (*6–8*). However, reducing the dosage to avoid over-silencing necessitates more frequent intrathecal administration, which in turn increases the risk of adverse events such as infection, cerebrospinal fluid leakage, neurovascular injury, and nerve or spinal cord damage (*9–12*). The variability in MECP2 expression levels among patients further complicates dosing strategies, necessitating individualized treatment regimens (*13*).

The MECP2 gene comprises four exons and three introns, with alternative splicing giving rise to two primary isoforms, MECP2 E1 and MECP2 E2. Despite differences in their N-terminal regions due to alternative splicing, both isoforms share identical DNA-binding and functional domains. MECP2 E1 comprises the majority of MECP2 in neurons and has a lower DNA-binding affinity compared to MECP2 E2 (*14*). The two isoforms are also expressed differentially depending on the stage of development, region and cell type of the brain (*15*). Isoform specific deletion studies have shown that MECP2 E1 ablation is sufficient to cause RTT like symptoms in mice while MECP2 E2 ablation was well tolerated in the context of RTT(*16, 17*). These observations, together with the fact that MECP2 E1 mutations have been found in RTT patients while MECP2 E2 mutations have not been seen in the clinic (*18–20*) provide further evidence that MEPC2 E1 loss-of-function is the main driver of the RTT phenotype. However, transgenic expression studies have demonstrated that either MECP2 E1 or MECP2 E2 can rescue the lethal phenotype and behavioral abnormalities observed in MECP2-null mice with the extent of behavioral and lifespan rescue related to expression levels of MECP2 regardless of isoform identity (*15, 21–24*). Similarly detailed information regarding the role of MECP2 isoforms is lacking for MDS. However, this functional redundancy suggests that in the context of MDS, selective silencing of one isoform could reduce the overall MECP2 expression level while preserving therapeutic efficacy and avoiding the need for dose titration.

A potential explanation for the two seemingly contradictory observations i.e. that i) MECP2 E1 deletion is sufficient to cause RTT and ii) RTT can be rescued by transgenic expression of MECP2 E2, is that natural levels of MECP2 E1 and the translation efficiency of E1 mRNA are greater than that of MECP2 E2, resulting in 90% of MECP2 expression in the brain being the E1 isoform (*25*). Therefore, studies showing that E1 ablation alone is sufficient to cause RTT may be confounded by differences in natural expression levels between the two isoforms, leading to the potentially erroneous conclusion that the two isoforms are not functionally redundant. In the context of MDS, the roles and contributions of each isoform has not been studied, therefore it is not known whether MECP2 E1 or MECP2 E2 alone can result in MDS when overexpressed.

An emerging class of NATs particularly well-suited to addressing the challenge of tunable yet durable silencing of MECP2 involves fully chemically modified siRNAs. These duplex RNA-like molecules are engineered to harness the endogenous microRNA-mediated gene silencing machinery, known as the RNA-induced silencing complex (RISC), to selectively bind to and degrade the target mRNA of disease-causing genes. These siRNAs are chemically modified to enhance plasma and tissue nuclease resistance by substituting every 2’ ribose position with 2’-fluoro or 2’-O-methyl groups, replacing the guide strand 5’-phosphate with 5’-vinylphosphonate, and protecting the 5’ and 3’ termini of both strands with phosphorothioate linkages instead of the naturally occurring phosphodiester backbone (*26–28*). siRNA-mediated gene silencing is catalytic, allowing a single siRNA-loaded RISC to bind and cleave multiple target mRNA molecules in the cytoplasm sequentially (*29*). We showed previously that divalent, fully chemically modified asymmetric siRNAs achieve broad central nervous system (CNS) distribution in mice and non-human primates (NHPs), enabling sustained gene silencing with a wide therapeutic index (*6*). Increased CNS retention and distribution likely stem from reduced CSF clearance, while phosphorothioate single-stranded regions may enhance cellular uptake efficiency (*28*).

The chemical modification of the siRNA scaffold provides a second approach to optimizing gene silencing. Due to extensive protein-RNA contacts and structural constraints within the siRNA-loaded RISC complex, 2’ ribose modifications must be strategically placed across the siRNA backbone to ensure that the local and global structure of the modified siRNA remains compatible with functional RISC loading. The specific positions of these chemical modifications—such as 2’-fluoro, 2’-O-methyl, and phosphorothioates—collectively constitute the modification pattern of the siRNA scaffold (*27*). While certain 2’-ribose modification positions have traditionally been viewed as incompatible due to their negative impact on siRNA efficacy(*30–32*), these positions can be reinterpreted as tunable sites. By strategically placing modifications at these sites, it is possible to fine-tune the efficacy of RISC loading and target mRNA cleavage, thus achieving a more controlled and durable gene silencing effect.

In this study, we developed fully chemically modified divalent siRNA therapeutic candidates capable of silencing total MECP2 or MECP2 E1. We evaluated these siRNA sequences in wild-type mice using two previously established 2’-ribose modification patterns, which have been shown to modulate siRNA efficacy across multiple sequences (*32–34*). In humanized transgenic mouse models of MECP2 duplication syndrome (MDS)(*35*), we demonstrated that excessive silencing of MECP2 in Tg1 mice (which expresses twice the endogenous MeCP2 levels) leads to mixed behavioral outcomes that could indicate toxicity, while moderate silencing does not produce the same effects. In the severe Tg3 MDS model (which expresses 3 to 5-fold the endogenous MeCP2 levels), isoform-selective silencing produced initial improvements in phenotypes but failed to achieve long-term disease modification with a single administration, while total MECP2 silencing modified disease for at least one year following a single administration. The identified pre-clinical candidates are cross-reactive to seven potential preclinical species, paving the way for formal pre-clinical development.

## RESULTS

### In vitro screen identifies potent multi-species targeting lead siRNAs

A comprehensive bioinformatic analysis of MECP2 transcripts, including the 5’ and 3’ untranslated regions (UTRs) and the open reading frame (ORF), was conducted to identify potential active siRNA sequences, following established guidelines(*36, 37*). Twenty-four top-scoring sequences, including those human-mouse cross-reactive sites, were synthesized for in vitro screening. Given that chemical modifications of siRNAs can significantly restrict the RISC-compatible sequence space, all siRNAs were synthesized using fully chemically modified, asymmetric siRNA scaffolds with 2’-fluoro, 2’-O-methyl, and phosphorothioate modifications, as previously described(*38*). To facilitate gymnotic uptake into cells without transfection reagents, the 3’ terminus of the passenger strand was covalently linked to cholesterol. Details of all sequences and chemical modification patterns used in this study are provided in the supplementary material (Table S1).

siRNA candidates targeting all MECP2 mRNA isoforms were screened in vitro at a single concentration of 1.5μM in human HeLa cells (Figure 1A) and mouse N2A cells (Supplemental Figure S1A), with gene expression assessed using the Quantigene 2.0 branched DNA assay. A non-targeting control siRNA (NTC) served as a negative control in all in vitro screens. siRNAs 1759 and 1764 exhibited significant *MECP2* silencing and were selected for further potency evaluation using a seven-point dose-response curve in both HeLa cells (Figure 1B) and N2A cells (Supplemental Figure S1B). The IC_50_ values for siRNA_1759 and siRNA_1764 were 201nM and 119nM, respectively, in passive uptake in HeLa cells, with comparable potency observed in N2A cells. These results indicate that these siRNAs are suitable candidates for further *in vivo* evaluation.

**Figure. 1.**
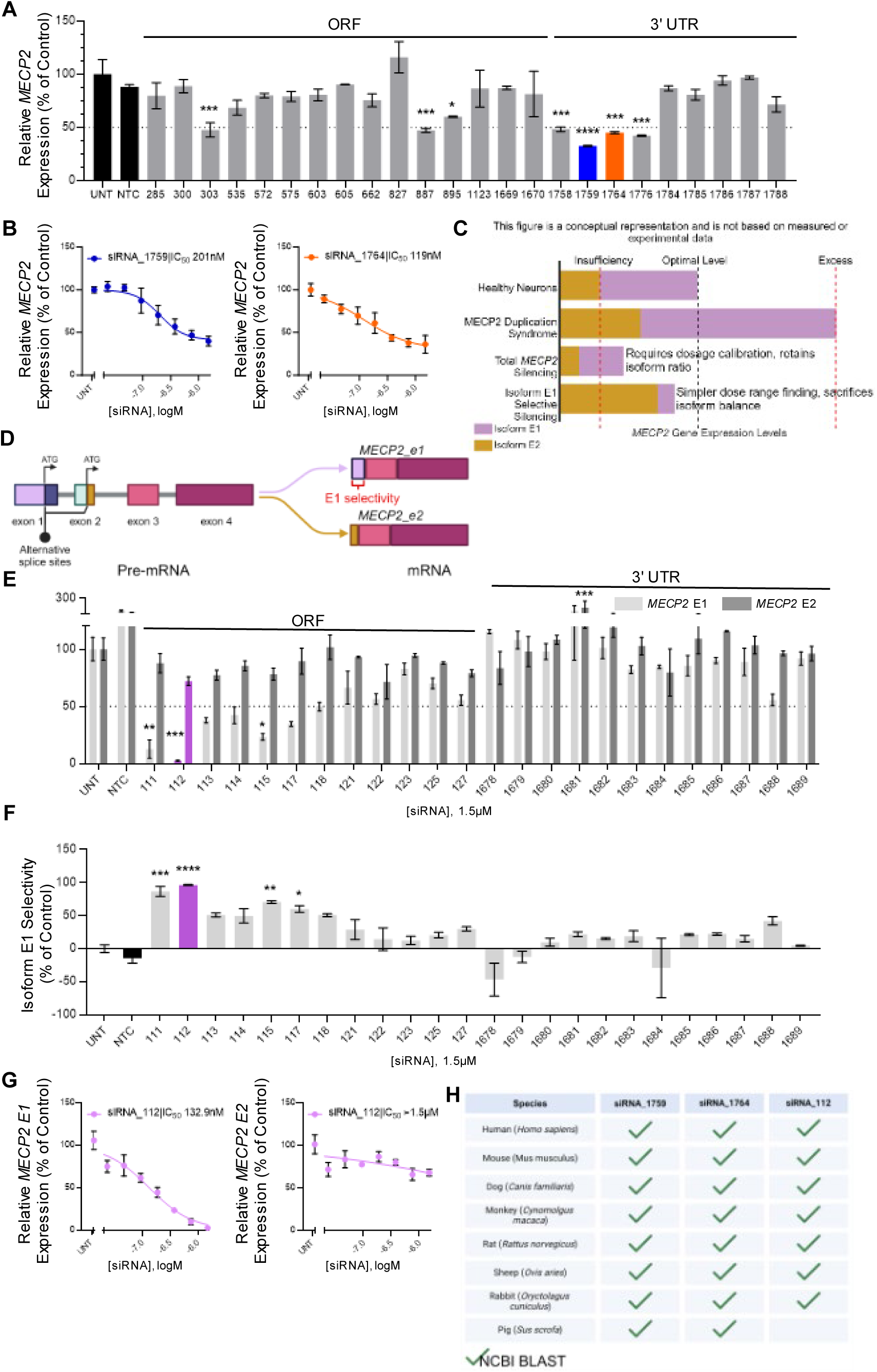
Lead total and isoform selective MECP2 silencing siRNAs identified through *in vitro* screens. (A) Relative expression of total human *MECP2* mRNA in HeLa cells 72h after treatment with non-isoform selective *MECP2* targeting siRNA at a single concentration. Lead compounds selected for further analysis are shown as blue (1759) and orange (1764) bars. (B) Relative expression of total human *MECP2* mRNA in HeLa cells 72h after treatment with total isoform silencing lead siRNAs 1759 (blue) and 1764 (orange) across a range of concentrations. (C) Potential strategies for modulation of MECP2 without dosage related toxicity. (D) Schematic of alternative splicing of the MECP2 pre-mRNA. (E) Relative expression of *MECP2-E1* (light grey bars) and *MECP2-E2* (dark grey bars) mRNA in HeLa cells 72h after treatment with E1 isoform selective siRNA at a single concentration. Lead compound (112) selected for further analysis is shown in purple bars. (F) Isoform E1 selectivity of siRNA compounds screened in (E). (G) Relative expression of *MECP2-E1* mRNA in HeLa cells 72h after treatment with E1 isoform selective siRNA 112 across a range of concentrations. (H) Results of cross-reactivity analysis of the targeting regions of lead siRNAs across the genome of humans and pre-clinical species using Basic Local Alignment Search Tool (BLAST) from the National Center for Biotechnology Information (NCBI) (green checkmarks) database. Gene expression was measured using the QuantiGene™ Singleplex Assay Kit. Data represented as mean±s.e.m. of three independent replicates. Statistical analysis performed using Ordinary one-way ANOVA with Dunnett’s adjustment for multiple comparisons (*-p<0.05, **-p<0.01, ***-p<0.001, ****-p<0.0001).

Given that MECP2 is a dosage-sensitive gene, where excessive silencing may lead to the development of Rett syndrome, fine-tuning the extent of silencing is critical for therapeutic development for MECP2 duplication syndrome (MDS)(*2*). While dosage adjustments can modulate silencing, lower doses of RNAi therapeutics typically compromise durability. Therefore, we explored a strategy to achieve tunable silencing without sacrificing therapeutic durability (Figure 1C). The two major MECP2 isoforms, E1 and E2, may have sufficient functional overlap to provide redundancy(*21*). In healthy neurons, both isoforms are expressed at levels that maintain optimal total MECP2. In MECP2 duplication syndrome, where duplications typically span multiple genes, both E1 and E2 are overexpressed, resulting in elevated MECP2 levels. Our strategy focuses on selectively silencing the E1 isoform, which is uniformly expressed across all brain cell types and at higher levels than E2. We hypothesize that silencing E1 to maximal achievable levels, without compromising dosage or duration, could reduce total MECP2 levels to within the optimal range. Isoform E2, which would remain predominantly expressed, should preserve sufficient MECP2 function to maintain healthy brain physiology. While E2 levels in wildtype mice are insufficient to maintain healthy brain physiology in the absence of E1, the increased expression levels of E2 in MDS may be conducive to this strategy.

We designed a panel of human and human-mouse cross-reactive siRNAs targeting exon 1, which is present in isoform E1 but absent in E2 (Figure 1D). HeLa cells were treated with 1.5μM of cholesterol-conjugated siRNAs designed to target MECP2 E1 without affecting E2. Expression levels of isoforms E1 and E2 were then measured using custom isoform-specific probes with the Quantigene 2.0 branched DNA assay. siRNA_112 provided maximal silencing of E1 (Figure 1E), with a ∼95% selectivity for E1 over E2 (Figure 1F). We were unable to find any siRNAs that specifically silenced the E2 isoform without affecting E1 (Supplementary Figure S1C). This result was corroborated in seven-point dose-response assays, where siRNA_112 silenced MECP2 E1 mRNA with an IC50 of 133nM in HeLa cells (Figure 1G) and 263nM in N2A cells (Supplemental Figure S1D), with minimal impact on MECP2 E2 mRNA expression.

The lead siRNAs—1759, 1764, and 112—identified from the in vitro screens targeted regions of MECP2 mRNA that were 100% conserved across multiple species, including mouse, dog, cynomolgus monkey, rat, sheep, rabbit, and pig, as determined by the Basic Local Alignment Search Tool (BLAST) from the National Center for Biotechnology Information (NCBI). (Figure 1H). The cross-species homology of these lead compounds ensures that formal IND-enabling preclinical studies will not be limited by the availability of on-target pharmacology models.

### Lead MECP2-targeting siRNAs tune MECP2 levels in wildtype mice for up to four months

To evaluate the *in vivo* potency and duration of effect for lead siRNAs, we synthesized these siRNAs using a validated divalent brain delivery scaffold, which has been shown to enable deep brain distribution and multi-month silencing across various animal models (*6*). The divalent siRNAs, composed of two siRNA passenger strands linked by a tetraethylene glycol linker and hybridized to the guide strand, were synthesized in two distinct modification patterns: scaffold 1 (∼59% 2’-O-methyl content) and scaffold 2 (∼76% 2’-O-methyl content), both of which have been previously demonstrated to modulate silencing for other siRNA sequences (*31, 33, 34*). Given that most clinically approved siRNAs feature a higher 2’-O-methyl to 2’-fluoro ratio (*30, 39*), likely due to the increased nuclease stability of 2’-O-methyl (*40*), we aimed to determine whether the 2’-O-methyl-rich scaffold 2 would offer greater durability for MECP2 silencing in the brain. The guide strand’s 5’-phosphate was stabilized using vinylphosphonate (VP) for enhanced binding affinity to the Mid domain of Argonaute-2 and enhanced phosphatase stability, while the ends of both strands were modified with phosphorothioate linkages for enhanced nuclease stability (*30, 39, 41*) (Figure 2A).

**Figure. 2.**
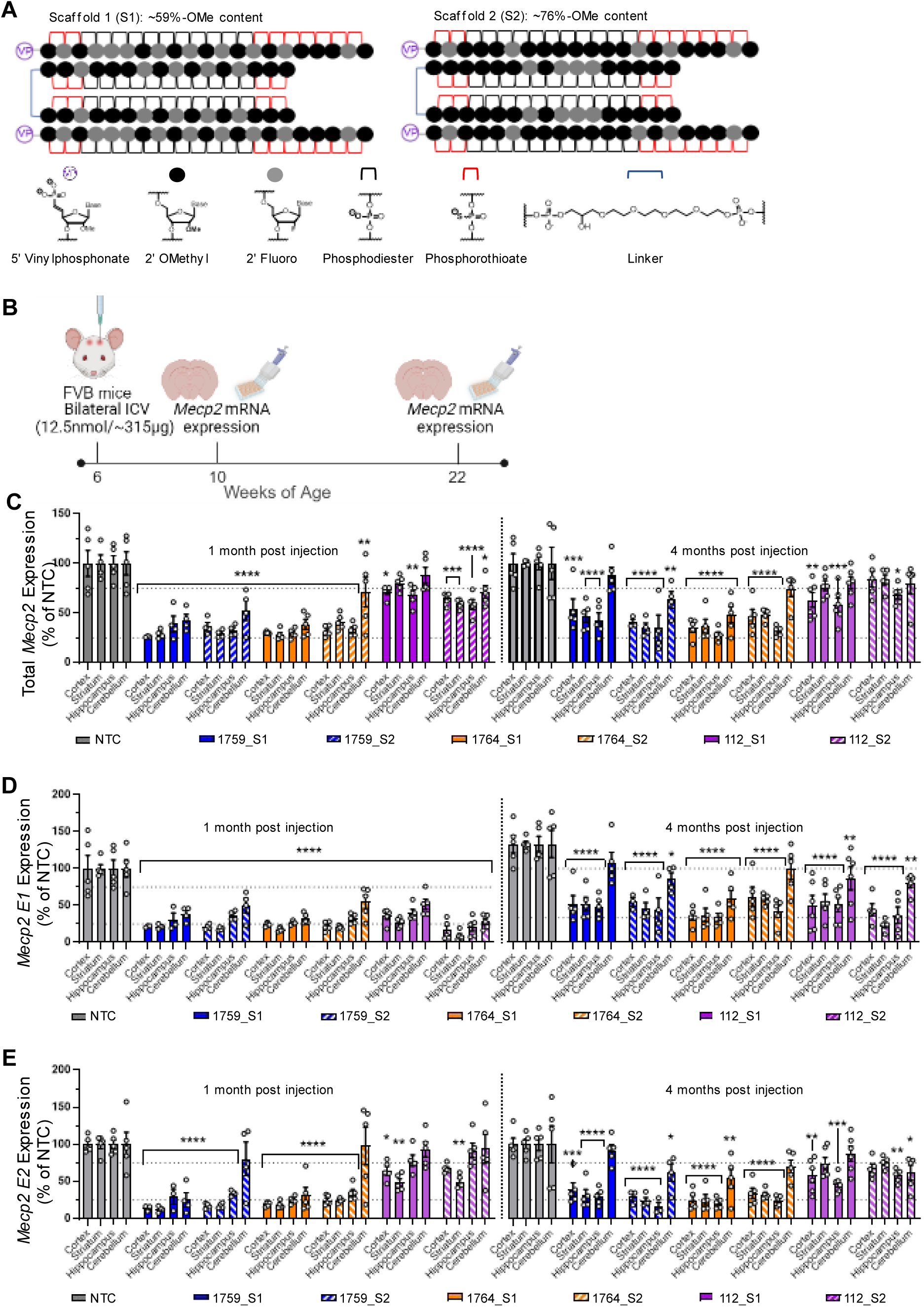
Lead compounds show potent and sustained modulation of *Mecp2* mRNA expression in the wildtype mouse brain. (A) Schematic of divalent scaffolds, modification patterns and chemical structures of relevant modifications used in this experiment. (B) Illustration of study design including dosage, route of administration, timeline and readouts. (C) Total *Mecp2* mRNA expression levels, (D) *Mecp2-E1* mRNA expression levels and (E) *Mecp2-E2* mRNA expression levels in the cortex, striatum, hippocampus and cerebellum of wildtype FVB mice injected with 12.5nmol (∼315 µg) of siRNA variants. Gene expression was measured at one month and four months post injection using the QuantiGene™ Singleplex Assay Kit. Data represented as mean±s.e.m. of individual animals (n=4-6 mice/group). Statistical analysis performed using two-way ANOVA with Dunnett’s adjustment for multiple comparisons (*-p<0.05, **-p<0.01, ***-p<0.001, ****-p<0.0001).

Female wildtype FVB mice were administered 12.5 nmol (∼315 µg) of divalent siRNAs or divalent non-targeting control (NTC) siRNA via intracerebroventricular (ICV) injection at six weeks of age. Mice were sacrificed at one- and four-months post-injection, and total *Mecp2*, *Mecp2-E1*, and *Mecp2-E2* mRNA expression levels were analyzed in the cortex, striatum, hippocampus, and cerebellum using the Quantigene 2.0 branched DNA assay (Figure 2B). While the Quantigene probesets for detecting E1 and E2 isoforms were custom designed for selectivity, the total *Mecp2* probeset detects a broader range of *Mecp2* mRNAs. Consequently, in this assay, the sum of E1 and E2 isoform expression cannot be used to infer total *Mecp2* mRNA levels. Instead, the E1 and E2 probesets were employed to assess siRNA selectivity, while the total *Mecp2* probeset served as the primary metric of clinical utility.

At one-month post-injection, total *Mecp2* mRNA levels in the cortex, striatum, and hippocampus of mice treated with non-isoform-selective siRNAs (1759_S1, 1759_S2, 1764_S1, and 1764_S2) were reduced to ∼25-35% of NTC controls (p<0.0001), with no significant differences observed across these brain regions (Figure 2C). Silencing in the cerebellum was less pronounced and more variable, due to lower siRNA accumulation in this region of rodent brains (*6*). In the cerebellum, mice treated with 1759_S1 and 1764_S1 showed mean total *Mecp2* expression levels of 43% (p<0.0001) and 38% (p<0.0001), respectively. In contrast, the 2’-O-methyl-rich scaffold 2 siRNAs (1759_S2 and 1764_S2) resulted in higher *Mecp2* levels of 53% (p<0.0001) and 71% (p<0.01), respectively (Figure 2C).

At four months post-injection, 1764_S1 retained near-maximal silencing in all brain regions tested, with total *Mecp2* mRNA levels between 35%-48% (p<0.0001) across all brain regions tested, making it the most potent compound for total *Mecp2* mRNA silencing in this study. In contrast, while 1759_S1, 1759_S2, and 1764_S2 maintained *Mecp2* mRNA levels at ∼35-45% of NTC controls in the cortex, striatum, and hippocampus (p<0.001 to p<0.0001), cerebellar *Mecp2* mRNA levels were 88% (p=0.7), 63% (p<0.01), and 74% (p<0.07), respectively, indicating reduced durability compared to 1764_S1 (Figure 2C).

For isoform E1-selective siRNAs (112_S1 and 112_S2), total *Mecp2* mRNA levels were reduced to 59%-72% across brain regions with this effect being maintained for at least four months. While 112_S2 showed better performance at one month, 112_S1 demonstrated superior durability for the duration tested (Figure 2C).

Isoform E1 mRNA levels in siRNA-treated mice at one-month post-injection were ∼16%-50% of NTC across all brain regions for all compounds tested (p<0.0001 for all groups). These results indicate that 112_S2 was the most potent suppressor of *Mecp2-E1* transcription in one month. Differences in efficacy between scaffolds S1 and S2 for 1759 and 1764 were minimal. Differences between scaffolds were more pronounced in the cerebellum, confirming that scaffold S2 was less active than S1 for 1759 and 1764 siRNAs, while 112_S2 remained the most effective at silencing E1 mRNA in the cerebellum (Figure 2D).

At four months post-injection, E1 mRNA levels across brain regions and compounds were 17%-44%. Although 112_S2 showed greater silencing at one month, 1764_S1 exhibited slightly, though not statistically significant, better durability in the cortex and hippocampus. This difference was more evident in the cerebellum, indicating that 1764_S1 provided the most durable silencing of *Mecp2-E1* (Figure 2D).

At one-month post-injection, suppression of *Mecp2* isoform E2 mRNA was notably higher than that of E1 or total *Mecp2* for the non-isoform-selective siRNAs (<25% of NTC). E1-selective siRNAs 112_S1 and 112_S2 also reduced E2 expression levels in the cortex and striatum but not the hippocampus and cerebellum, indicating that these siRNAs did not achieve full E1 isoform selectivity at the tested doses in certain brain regions. Similar to total and E1 isoform *Mecp2* mRNA levels, the lower accumulation of divalent siRNA in the rodent cerebellum allowed for clearer differentiation between scaffolds S1 and S2, with scaffold S2 showing lower efficacy in this region (Figure 2E).

At four months post-injection, 112_S1 and 112_S2 had some impact on E2 mRNA levels, but this effect did not reach statistical significance except in the hippocampus. The effect of non-selective 1759 and 1764 were similar to that observed at the one month time point. In the cerebellum, the differences between siRNAs were more pronounced, with no significant silencing observed in mice treated with 1759_S1, 112_S1, or 1764_S2 (Figure 2E).

In general, all non-selective compounds maintain robust silencing of total *Mecp2* with less than 30-40% of total mRNA remaining at up to four-months post injection. There were no significant differences between scaffolds except for 1764_S1 showing higher potency in the cerebellum compared to 1764_S2. Based on this data, 1764 S1 was selected for further evaluation in disease models. The isoform selective compound 112_S1 was selected as a second lead compound to evaluate the effects of E1 silencing with minimal perturbation of E2 levels. We have previously seen differences between protein and mRNA silencing for CNS targets due to the presence of inaccessible nuclear pools of target mRNA (*42*). Indeed, we later evaluated the MECP2 silencing in the context of the animal models and confirmed that ∼ 60-70% mRNA silencing corresponds to highly potent protein reduction of ∼ 80-90% (Figure 3).

**Figure. 3.**
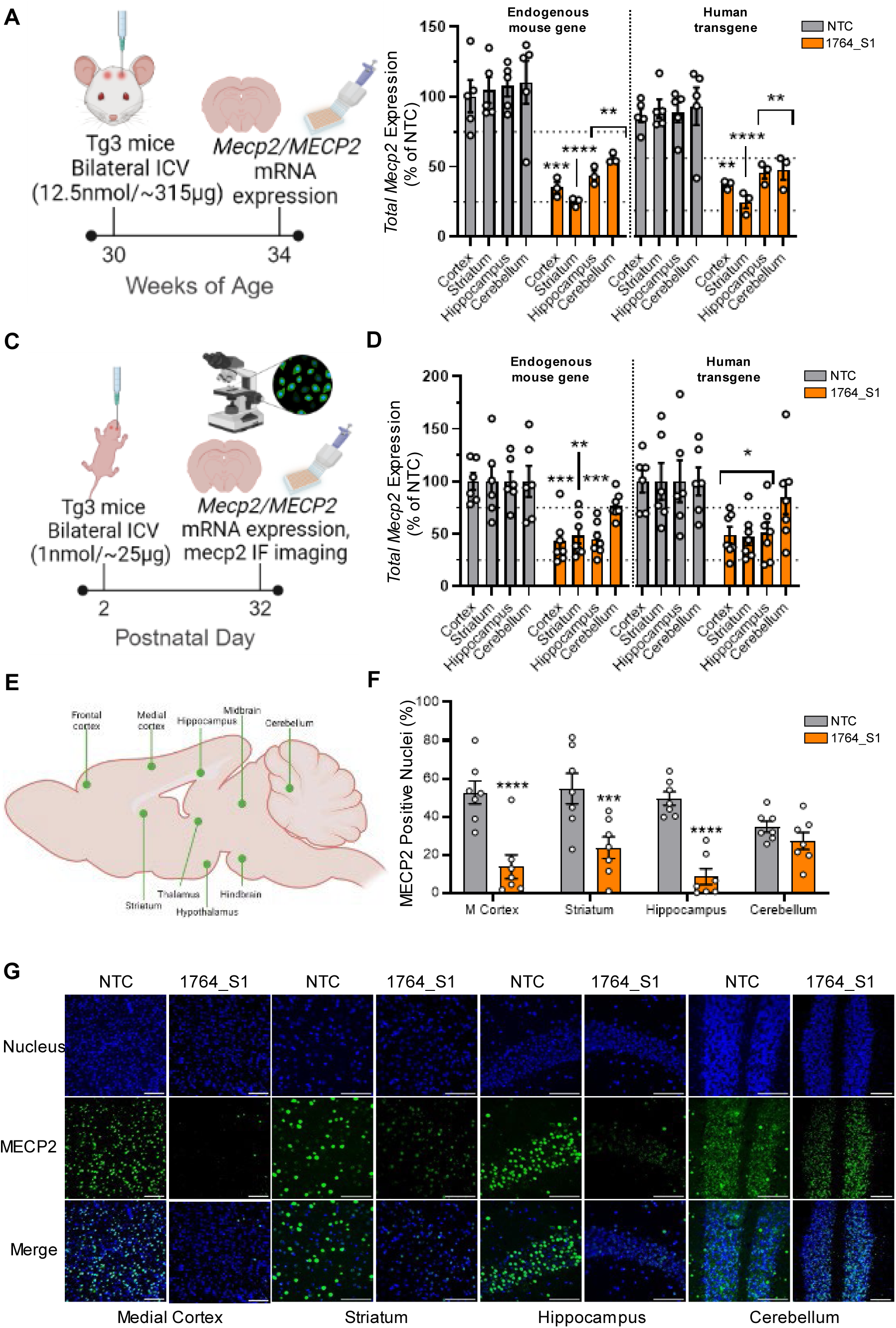
1764_S1 is active against mouse and human MECP2 in the adult and neonatal Tg3 brain. (A) Illustration of study design including dosage, route of administration, timeline and readouts in adult Tg3 mice. (B) Total mouse *Mecp2* and human mouse *MECP2* mRNA expression levels in the cortex, striatum, hippocampus and cerebellum of adult Tg3 mice injected with 12.5nmol (∼315 µg) of 1764_S1 (n=5 mice for NTC and n=3 mice for 1764_S1). (C) Illustration of study design including dosage, route of administration, timeline and readouts in neonatal Tg3 mice. (D) Total mouse *Mecp2* and human mouse *MECP2* mRNA expression levels in the cortex, striatum, hippocampus and cerebellum of neonatal Tg3 mice injected with 1nmol (∼25 µg) of 1764_S1 (n=6 for NTC and n=7 for 1764_S1). (E) Illustration of brain regions of interest identified for relative quantification of immunofluorescence from mice shown in (C). (F) Percentage of MECP2 positive nuclei in the medial cortex (M. cortex), striatum, hippocampus and cerebellum of neonatal mice shown in (C). (G) Representative immunofluorescence images at 40x magnification of samples quantified in (F), scale bar = 100µm. Gene expression was measured using the QuantiGene™ Singleplex Assay Kit. Data represented as mean±s.e.m. of individual animals. Immunofluorescence imaging was performed on sagittal brain sections stained with DAPI and anti-MECP2 antibody and imaged using Leica DMi8 widefield microscope. Quantification of MECP2 positive nuclei was performed using a custom imageJ script. Statistical analysis performed using two-way ANOVA with Dunnett’s adjustment for multiple comparisons (*-p<0.05, **-p<0.01, ***-p<0.001, ****-p<0.0001).

An interesting finding that warrants further exploration (see discussion) is the absence of overt toxicity phenotypes when MECP2 protein levels were reduced at least 70-80% throughout the wild-type mouse brain. However, it must be noted that this study was not designed to look carefully into potential behavioral phenotypes, and studies have shown that even reductions in MECP2 levels of 20-30% in wildtype mice result in anxiety-like phenotypes and other behavioral deficits that mimic some of the phenotypes of MECP2-null mice (*43*).

### Activity of 1764_S1 against human MECP2 confirmed in a severe transgenic mouse model of MDS

To account for the variability in duplication size and MECP2 overexpression among MDS patients, we selected two well-characterized human transgenic MDS models developed by the Zoghbi lab (*35*)—MECP2Tg1 (Tg1), which expresses MECP2 at approximately two-fold the wild-type level, and MECP2Tg3 (Tg3), which expresses MECP2 at approximately three-to five-fold the wild-type level—to evaluate the efficacy of the lead siRNA. These transgenic lines were created by microinjection of a 99kb human PAC clone carrying the entire MECP2 locus, ensuring expression of both isoforms including with untranslated regulatory regions. Initially, we aimed to verify the in vivo efficacy of 1764_S1 for silencing human MECP2 in the Tg3 transgenic line before proceeding to test both lead siRNAs for their potential to modulate MDS in Tg1 and Tg3 models, thereby prioritizing animal ethics and resource efficiency.

Adult Tg3 mice at 30 weeks of age were injected with 12.5 nmol (∼315 µg) of 1764_S1 via bilateral intracerebroventricular (ICV) injection. The mice were sacrificed at 34 weeks of age, and their brains were analyzed for total MECP2 mRNA levels (Figure 3A). Since these transgenic mice are hemizygous, containing both the endogenous mouse gene and the human transgene, we were able to measure the total MECP2 mRNA of both species using species-specific probesets in the QuantiGene™ Singleplex branched DNA assay. As expected, total mouse *Mecp2* mRNA levels in the cortex, striatum, hippocampus, and cerebellum were 35% (p < 0.001), 25% (p < 0.0001), 44% (p < 0.01), and 56% (p < 0.01) of non-targeting control (NTC) levels, respectively. Silencing of total human *MECP2* mRNA was slightly lower, with levels of 50% (p < 0.01), 33% (p < 0.0001), 61% (p < 0.01), and 64% (p < 0.01) in the cortex, striatum, hippocampus, and cerebellum, respectively. These levels across all four brain regions were close to ∼50% of NTC controls, which we consider a tolerable yet disease-modifying level of efficacy (Figure 3B).

Given that MDS is a pediatric condition where early intervention is presumably more beneficial, we also assessed whether safe levels of silencing could be achieved in the smaller brains of neonatal mice. Tg3 mice at postnatal day 2 were injected with 1 nmol (∼25 µg) of 1764_S1 via bilateral ICV injection, and the mice were sacrificed at postnatal day 32. Subsequently, total MECP2 human and mouse mRNA levels were measured using the QuantiGene™ Singleplex branched DNA assay, and total MECP2 protein levels were analyzed by immunofluorescence in different brain regions (Figure 3C). It should be noted that due to the small size of the neonatal mouse brain, the ICV injections were performed manually, leading to potential variability in the speed of injection and distribution of the siRNA throughout the brain. Total mouse *Mecp2* mRNA levels in the cortex, striatum, hippocampus, and cerebellum were 43% (p < 0.001), 48% (p < 0.01), 45% (p < 0.001), and 77% (p = 0.3) of NTC controls, respectively. Total human *MECP2* mRNA levels in the cortex, striatum, hippocampus, and cerebellum were 49% (p < 0.01), 47% (p < 0.01), 51% (p < 0.01), and 85% (p = 0.9) of NTC controls, respectively (Figure 3D).

For immunofluorescence quantification, whole brain tiled scans were collected at 40x magnification. Specific brain regions—namely the frontal cortex, striatum, medial cortex, hippocampus, thalamus, hypothalamus, midbrain, hindbrain, and cerebellum—were identified and cropped for analysis (Figure 3E). Image thresholding was used to accurately separate the DAPI (nucleus) fluorescence signal from the background, followed by quantification using the ImageJ Measure Function. Individual nuclei were identified and counted as regions of interest (ROIs), and within each ROI, MECP2-positive nuclei were counted and expressed as a percentage of the total nuclei in each brain region. On average, 500 to 1,000 nuclei were identified for each brain region per animal.

MECP2 protein positivity was significantly lower in 1764_S1-treated mice compared to NTC controls across most regions measured. The percentage of MECP2-positive nuclei for each region, expressed as 1764_S1 vs. NTC, was as follows: 10% vs. 50% (p < 0.0001) in the prefrontal cortex, 24% vs. 55% (p < 0.001) in the striatum, 14% vs. 53% (p < 0.0001) in the medial cortex, 8% vs. 50% (p < 0.0001) in the hippocampus, 18% vs. 47% (p < 0.01) in the thalamus, 26% vs. 52% (p < 0.01) in the hypothalamus, 35% vs. 50% (p = 0.3) in the midbrain, 30% vs. 49% (p = 0.1) in the hindbrain, and 27% vs. 35% (p = 0.9) in the cerebellum (Figure 1F and Supplemental Figure S2F). Representative immunofluorescence images of each brain region from NTC and 1764_S1-treated mice are shown in Figure 3G and Supplemental Figures S2A-E. As observed in adult mice, 1764_S1 effectively reduced total MECP2 mRNA and protein levels in multiple brain regions of Tg3 neonatal mice.

Interestingly, the level of protein silencing observed was much greater than the levels of mRNA silencing, indicating that there may be a fraction of transcribed mRNA retained in the nucleus and inaccessible to cytoplasmic siRNA activity, as has been reported for huntingtin, another CNS target (*42*). More importantly, it underscores the importance of the MECP2 protein level as the primary measure of pharmacodynamics since quantification of the mRNA alone may underestimate the degree of silencing. Further work on dose titration using MECP2 protein levels as a readout of pharmacological activity is essential to the preclinical development of these siRNAs.

### Single injection of non-isoform selective 1764_S1 but not E1 selective 112_S1 induces mixed behavioral effects in Tg1 mice

To evaluate the efficacy of the lead compounds in modifying disease and the potential for Rett-like toxicity due to excessive suppression, we utilized the mild Tg1 model of MDS. This model is known to overexpress the human MECP2 transgene at approximately twice the levels observed in wild-type FVB mice. The Tg1 mice were obtained from Jackson Laboratory (JAX) via Speed Expansion, using cryopreserved sperm from Tg1 males and oocytes from wild-type FVB females for the first cohort, and subsequent cohorts were produced by mating mutant females with wild-type (WT) males. MECP2 overexpression was confirmed by JAX prior to the study, with Western blot data indicating an ∼2.4-fold increase in MECP2 expression compared to FVB wild-type controls (Supplemental Figure S3A). Behavioral assays were chosen based on published data from MDS transgenic mouse models and were conducted using standard operating procedures established by JAX In Vivo Pharmacology Services. The study was performed by JAX.

Adult six-week-old male Tg1 mice were administered 12.5 nmol (∼315 µg) of either 1764_S1, 112_S1, or phosphate-buffered saline (PBS) as a vehicle control via intracerebroventricular (ICV) injection. We elected to use a single PBS control group as we have previously established the equivalency of NTC and PBS on target silencing *in vitro* (see Fig. 1A, E and (see F) and *in vivo* (*6*). Wild-type littermates received PBS as a negative control for the MDS genotype. Technicians were blinded to the content of each test article throughout the study. Post-administration, the mice were monitored for up to a year before euthanasia. Behavioral assessments included nest building at 15 and 25 weeks of age, the Open Field Test at 16 and 26 weeks of age, and rotarod performance at 17 and 27 weeks of age (Figure 4A). These timepoints were selected to evaluate behavior at a reasonable onset age and to track disease progression based on previous publications (*35*). The experiment aimed to assess the potential of lead siRNAs to alter behavior and mortality in MDS mice; tissue samples were not collected for molecular biology analyses before euthanasia.

**Figure 4.**
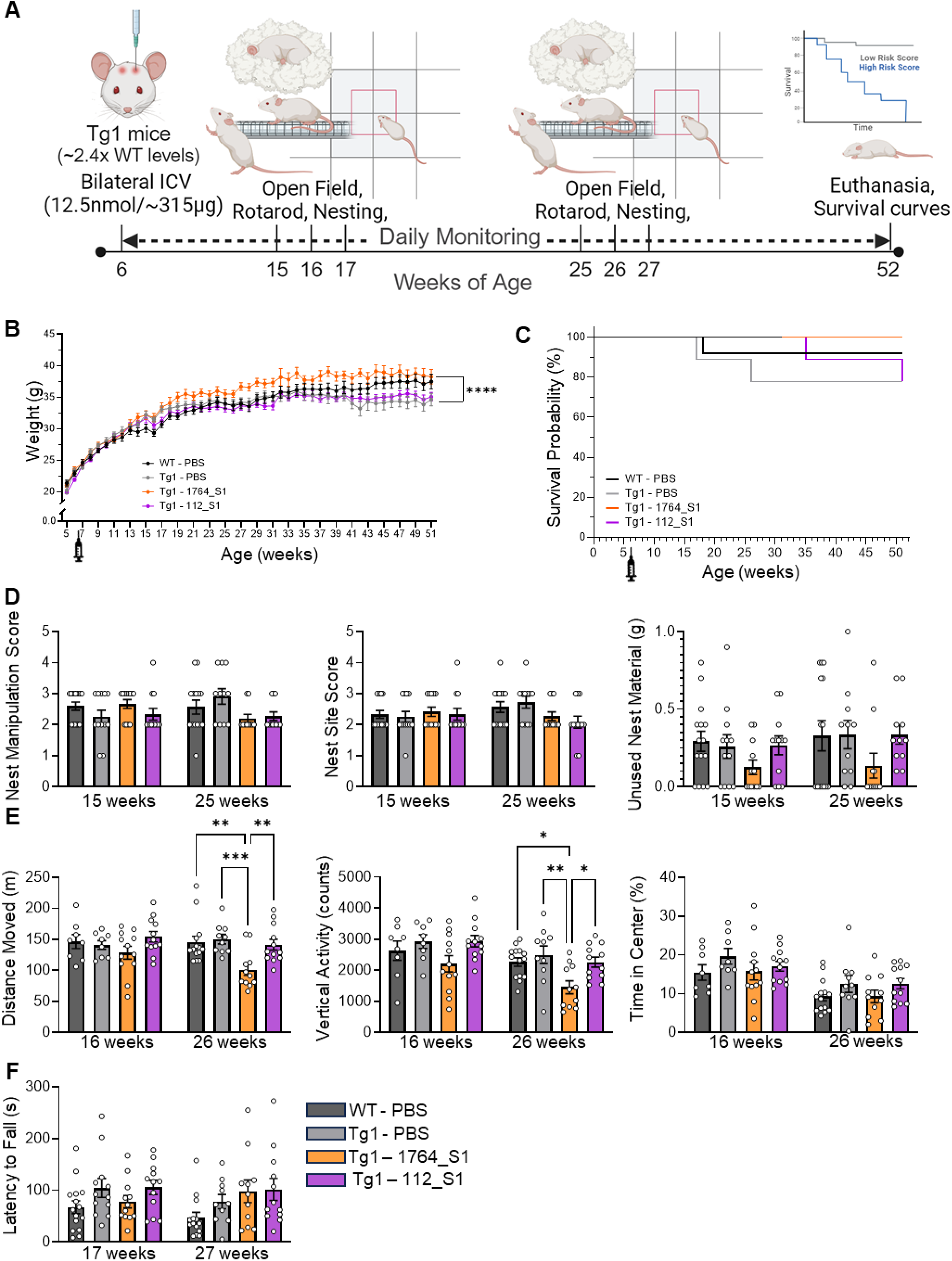
1764_S1 but not 112_S1 induces mixed behavioral effects in Tg1 mice. (A) Illustration of study design including dosage, route of administration, timeline and readouts in WT and Tg1 mice treated with PBS (black data points – WT, grey data points – Tg1), 1764_S1 (orange data points) and 112_S1 (purple data points). (B) Weight of treated animals over time. (C) Mortality of treated animals over time expressed as the probability of survival. (D) Nest building assay scores including nest manipulation score (left), nest site score (middle) and unused nesting material (right) measured at 15 and 25 weeks of age. (E) Open field test results including distance moved (left), vertical activity counts (middle) and time in the center (right) measured at 16 and 26 weeks of age. (F) Latency to fall on the rotarod test measured at 17 and 27 weeks of age. Data represented as mean±s.e.m. of individual animals (n=12-15 mice per group). Statistical analysis performed using two-way ANOVA with Dunnett’s adjustment for multiple comparisons (**-p<0.01, ****-p<0.0001).

The body weight of Tg1 mice administered PBS was comparable to that of wild-type FVB (WT) mice given PBS, indicating no genotype effects on weight or weight gain in this experiment. Tg1 mice treated with the E1 isoform-selective 112_S1 also showed similar weights to Tg1 PBS and WT PBS mice. Interestingly, Tg1 mice treated with the non-isoform-selective 1764_S1 exhibited a statistically significant weight gain compared to Tg1 PBS controls, with the two groups diverging in weight after 16 weeks of age (Figure 4B). Survival analysis revealed no significant differences in survival up to 52 weeks of age for either WT or Tg1 mice, and no treatment effects were observed on mortality (Figure 4C). The reason for this discrepancy in mortality compared to previous reports of mortality from 30 weeks up to a year of age (*35*) is unclear as protein analysis of WT and Tg1 brains at four weeks of age confirmed transgene overexpression. One potential explanation is that in this study, mice were separated and individually housed due to excessive fighting which may have prevented aggression related mortality. Increased aggression in male mice with ∼50% transgenic overexpression of Mecp2 in the FVB but not in the C57BL/6 background has been reported previously (*44*).

Nest building, previously reported as a marker of disease in Rett and MDS mouse models (*44, 45*), was assessed by providing each mouse with a single nestlet in their home cage. After 24 hours, nest photographs were scored based on nest manipulation and site quality, and the weight of unused nest material was recorded. At both 15 and 25 weeks of age, transgene overexpression did not significantly impact nest-building parameters and treatment with isoform selective 112_S1 did not significantly change nest building. However, treatment with non-isoform selective 1764_S1 resulted in non-significant decreases in the unused nest material suggesting over-manipulation of nests compared to wildtype and vehicle treated Tg1 mice. (Figure 4D). In the Open Field Test, conducted at 16 and 26 weeks of age, Tg1 PBS animals did not exhibit any change in behavior compared to wildtype controls. Treatment of Tg1 mice with isoform selective 112_S1 did not alter this behavior. However, treatment with the non-isoform selective 1764_S1 significantly reduced the distance moved and vertical activity at 26 weeks of age (Figure 4E). The rotarod test, administered at 17 and 27 weeks of age, showed no statistically significant differences in the latency to fall between any of the groups tested (Figure 4F).

Collectively, these results suggest that in our study, the Tg1 MDS mice did not experience severe disease, and showed no changes in survival probability or behavior relative to WT controls. However, treatment with the non-isoform-selective 1764_S1 did significantly increase body weight, decrease activity in the Open Field, and possibly increase manipulation of nest material in the nest building test. These behavioral differences in 1764_S1, but not 112_S1 treated Tg1 mice may indicate adverse effects due to exaggerated pharmacological activity of the siRNA. However, it remains uncertain whether 112_S1 treatment can safely modify disease in Tg1 mice due to the lack of an overt disease phenotype observed in this experiment.

### A single administration of siRNA 112_S1 partially and siRNA 1764_S1 completely modifies disease in severe Tg3 MDS mice

Given the lack of a clear MDS phenotype in Tg1 mice, we turned to a more severe MDS model, the Tg3 mice, to evaluate the effectiveness of MECP2 silencing siRNA in disease modification. This model reportedly overexpresses the human MECP2 transgene at approximately 3.5-fold higher levels than wild-type FVB mice. The Tg3 mice were obtained from Jackson Laboratory (JAX) through Speed Expansion, using cryopreserved sperm from Tg3 males and oocytes from wild-type FVB females for the first cohort with subsequent cohorts generated by mating mutant females with WT males. MECP2 overexpression was confirmed by JAX before the study, with Western blot data showing an ∼8.4-fold increase in MECP2 expression compared to FVB wild-type controls (Supplemental Figure S3B). Behavioral assays were conducted using standard operating procedures established by JAX In Vivo Pharmacology Services. The study was conducted at JAX.

Adult six-week-old male Tg3 mice were administered 12.5 nmol (∼315 µg) of either 1764_S1, 112_S1, or phosphate-buffered saline (PBS) as a vehicle control via intracerebroventricular (ICV) injection. Wild-type littermates received PBS as a negative control for the MDS genotype. Technicians were blinded to the contents of each test article throughout the study. The mice were monitored for up to 40 weeks post-administration before euthanasia. Behavioral assessments included nest building at 11 and 21 weeks of age, the Open Field Test at 12 and 22 weeks of age, and rotarod performance at 13 and 23 weeks of age (Figure 5A). These time points were chosen to evaluate behavior at a reasonable onset age and to track disease progression. The study aimed to assess the potential of lead siRNAs to alter behavior and mortality in MDS mice; tissue samples were not collected for molecular biology analyses before euthanasia.

**Figure 5.**
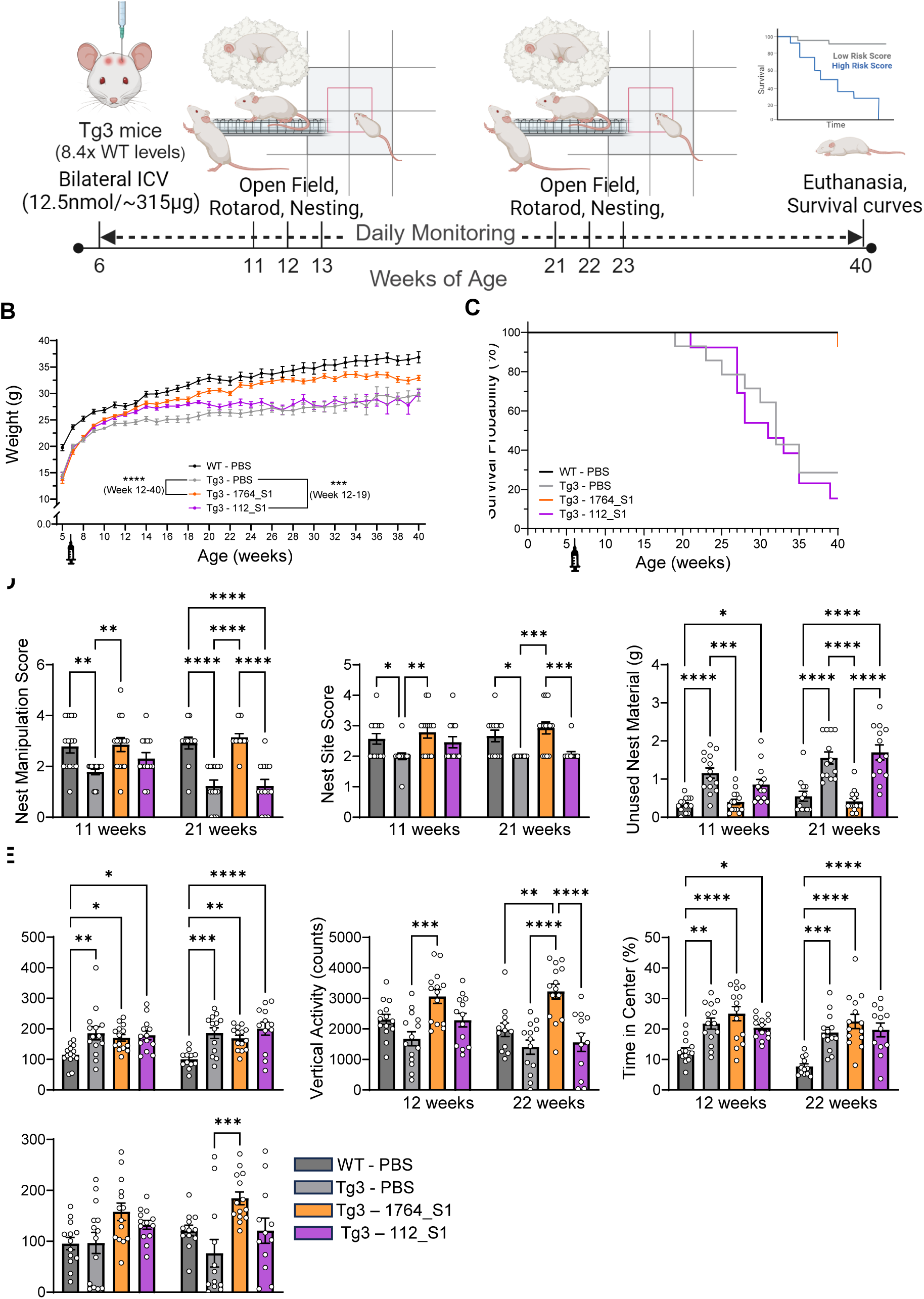
112_S1 partially and 1764_S1 completely rescues severe MDS in Tg3 mice. (A) Illustration of study design including dosage, route of administration, timeline and readouts in WT and Tg3 mice treated with PBS (black data points – WT, grey data points – Tg1), 1764_S1 (orange data points) and 112_S1 (purple data points). (B) Weight of treated animals over time. (C) Mortality of treated animals over time expressed as the probability of survival. (D) Nest building assay scores including nest manipulation score (left), nest site score (middle) and unused nesting material measured at 11 and 21 weeks of age. (E) Open field test results including distance moved (left), vertical activity counts (middle) and time in the center measured at 12 and 22 weeks of age. (F) Latency to fall on the rotarod test measured at 13 and 23 weeks of age. Data represented as mean±s.e.m. of individual animals (n=12-14 mice per group). Statistical analysis performed using two-way ANOVA with Dunnett’s adjustment for multiple comparisons (*-p<0.05, **-p<0.01, ***-p<0.001, ****-p<0.0001).

Tg3 male mice started at lower body weights than WT males. However, both the non-isoform-selective 1764_S1 and the E1 isoform-selective 112_S1 increased the body weight of Tg3 mice to WT levels, with statistically significant weight differences from PBS-treated Tg3 mice observed between 12-19 weeks of age for 112_S1-treated Tg3 mice (p<0.001). After 19 weeks of age, 112_S1-treated mice gained weight at a slower rate, aligning with Tg3 PBS levels thereafter. In contrast, 1764_S1-treated Tg3 mice showed significant weight improvement (p<0.0001), maintaining this gain throughout their lifespan (Figure 5B).

Unlike Tg1 males, Tg3 mice exhibited early mortality, with a median survival probability of 32 weeks. Treatment with the E1 isoform-selective 112_S1 did not affect early mortality, resulting in a median survival probability of 31 weeks. Notably, 1764_S1-treated mice did not exhibit any signs of mortality up to 40 weeks of age, the study’s endpoint, showing survival indistinguishable from WT PBS mice (Figure 5C).

The nest-building test, conducted at 11 and 21 weeks of age, was significantly impacted by genotype, with Tg3 PBS males showing lower nest manipulation scores, lower nest site scores, and higher amounts of unused nest material compared to WT PBS males at both time points. Treatment with the E1 isoform-selective 112_S1 slightly improved nest manipulation and site scores and reduced unused nest material at 11 weeks compared to Tg3 PBS mice, but these improvements were not statistically significant and were not sustained at 21 weeks of age. In contrast, Tg3 mice treated with 1764_S1 showed a complete rescue of nest manipulation and site scores and unused nest material at 11 weeks of age, maintaining these improvements at 21 weeks of age, with scores closely resembling those of WT PBS mice (Figure 5D).

The Open Field Test, administered at 12 and 22 weeks of age, revealed genotype effects, with Tg3 PBS males exhibiting increased distance moved, reduced vertical activity, and increased time spent in the center compared to WT PBS males. Neither 1764_S1 nor 112_S1 treatments rescued the increased distance moved or time spent in the center, as all Tg3 cohorts displayed similar patterns regardless of treatment. Although 112_S1 treatment slightly increased vertical activity to WT levels at 12 weeks of age, this effect was not statistically significant, and by 22 weeks of age, vertical activity in 112_S1-treated Tg3 mice aligned with Tg3 PBS controls. However, 1764_S1 treatment significantly increased vertical activity at both 12 weeks of age (p<0.001) and 22 weeks of age (p<0.0001) compared to Tg3 PBS controls, with activity levels even surpassing those of WT PBS males, indicating an overcorrection similar to that observed in Tg1 mice (Figure 5E).

The rotarod test, administered at 13 and 23 weeks of age, showed no genotype effects at 13 weeks, with similar latency to fall times for Tg3 PBS and WT PBS males. However, by 23 weeks of age, Tg3 PBS animals exhibited a reduced latency to fall compared to WT animals. Treatment with 112_S1 improved latency to fall in Tg3 animals at both 13 and 23 weeks, although this improvement was not statistically significant. In contrast, 1764_S1 treatment significantly increased latency to fall at 13 weeks of age (p<0.05) and 23 weeks of age (p<0.001) compared to Tg3 PBS males, with latency times even surpassing those of WT PBS males (Figure 5F).

It is noteworthy that, despite the early mortality observed in Tg3 PBS males, the survival probability of this genotype was significantly higher than in the original study describing these mice where premature death was seen as early as 2-4 weeks of age (*35*). This suggests that the MDS phenotype in both Tg1 and Tg3 lines has been significantly modified since the transgenic lines were first studied.

In conclusion, selective modulation of MECP2 improved phenotypes, though effects diminished after four months without impacting overall survival. Conversely, non-selective MECP2 modulation through a single treatment at six weeks of age led to significant neurobehavioral phenotype recovery and completely reversed the premature mortality observed in both control and isoform-selective treatment groups. These results indicate that treatment with isoform selective 112_S1 may be a viable option to avoid potential toxicity due to over silencing with 1764_S1, but repeat dosing may be necessary to maintain the beneficial effects of isoform selective silencing.

## DISCUSSION

The development of nucleic acid therapeutics for treating rare genetic diseases holds immense promise, offering potential cures for conditions that currently rely solely on symptomatic management. However, addressing diseases involving dosage-sensitive genes such as MECP2 presents significant challenges, as both over-correction and under-correction can lead to pathological outcomes. In the case of MECP2 Duplication Syndrome (MDS), excessive silencing of MECP2 may induce Rett-like syndromes, which are equally debilitating (*2*). Therefore, the preclinical development of nucleic acid therapeutics for MDS must ensure that MECP2 silencing is robust, tunable, and durable to achieve therapeutic efficacy without adverse effects. One innovative approach we employed involves isoform-selective silencing, leveraging the functional overlap between the two *MECP2* isoforms, E1 and E2, where one isoform can potentially compensate for the loss of the other (*21*). We developed small interfering RNAs (siRNAs) that target all isoforms of *MECP2* non-selectively for severe duplication cases, as well as siRNAs that selectively target the E1 isoform with minimal impact on E2 for milder cases where the risk of treatment-related Rett-like syndromes is highest.

A single dose of lead siRNAs achieved up to 80% MECP2 mRNA silencing at four months post-injection, with maximum silencing levels varying by isoform selectivity and chemical modifications. In the severe MDS model (Tg3), a non-isoform selective siRNA (1764_S1) fully rescued weight loss, behavioral deficits, and survival for the one-year study period, marking the first observed survival benefit in MDS therapy with a single dose. E1 isoform-selective silencing using 112_S1 provided initial gains in weight and behavioral trends in Tg3 but lost efficacy after 19 weeks, despite similar durability in wildtype mice. This suggests that single-dose E1-selective silencing may not be clinically viable but could prove effective in milder cases or with repeated dosing. In the milder MDS model (Tg1), which resembles clinical gene dosage, a disease phenotype was not apparent compared to wildtype littermate controls. In the absence of measurable disease, treatment with 1764_S1 resulted in changes in nest building behavior and rotarod performance. Mice treated with 1764_S1 also showed a statistically significant weight increase over Tg1 controls, though not significant compared to wildtype mice. These findings indicate potential 1764_S1 toxicity emerging over time in this mild disease model.

A major challenge in this study was the delayed onset of phenotypes in the Tg1 and Tg3 MDS models compared to published data. In Tg3 mice, median survival was 31 weeks, with the first death post-15 weeks, whereas literature reports mortality as early as 2-4 weeks. In the milder Tg1 model, no premature death or substantial behavioral phenotype was seen up to one year, unlike published accounts of seizures, hypoactivity, kyphosis, and death starting at 30 weeks (*35*). While we were unable to investigate the cause of this transgenerational phenotypic variation, a variety of factors including epistasis, compensatory mechanisms and environmental factors could be contributing to the observed dampening of MDS phenotypes in these transgenic lines (*46–48*).

Consistent with published reports, silencing durability varied by siRNA scaffold modifications, highlighting potential for tunable silencing in dosage-sensitive genes and pediatric cases (*30, 49*). A remarkable finding was the profound survival benefit and enduring phenotypic improvements observed for up to one year following a single administration of the 1764_S1 lead compound at six weeks of age. The study’s complexity precluded tissue collection at survival endpoints, resulting in a lack of information regarding the exact duration of effect. However, previously published reports using divalent siRNA and ICV delivery in the mouse brain have demonstrated at least six months of pharmacodynamic activity for similar doses (*6*). Given that the siRNA scaffold and not the sequence determines durability and pharmacokinetics, it is reasonable to conclude that the duration of silencing in this experiment was at least six months. However, due to the absence of silencing information, it remains uncertain whether the phenotypic benefits of 1764_S1 persisted beyond the pharmacodynamic effect, similar to reports with antisense oligonucleotides in MDS (*3*).

The siRNAs reported here are promising due to their extended effect duration compared to advanced antisense oligonucleotides like ION440, set for clinical trials by year’s end (Ionis Pharmaceuticals, ClinicalTrials.gov ID NCT06430385). Preclinical ION440 studies show MECP2 mRNA and protein silencing for 6-8 weeks following a single 500 µg dose (*3*). Since nucleic acid therapeutics generally cannot cross the blood-brain barrier (BBB) without a BBB shuttle or specific scaffold (*50–53*), intrathecal (IT) or intracerebroventricular (ICV) administration is required, posing significant patient risks (*9–12*). Therefore, durability is crucial for clinical utility and acceptance, particularly in the pediatric population. Additionally, ASOs accumulate primarily in the cortex, with lower levels of silencing in deeper brain regions like the putamen and the caudate compared to the more homogenous distribution and greater deep brain penetration of divalent siRNAs across preclinical species (*54–56*).

Clinical translation of these novel siRNAs will require further characterization of isoform-selective vs. non-selective silencing effects on the brain’s transcriptomic landscape, with comparisons to MECP2 knockout and overexpression controls via RNAseq. Preclinical toxicity studies must establish clear “no adverse effect levels” (NOAELs) in two species, ensuring safety at clinically effective doses. While MECP2 loss is linked to Rett syndrome, mainly studied in knockout models, these siRNAs will help determine the MECP2 reduction required to trigger Rett-like phenotypes postnatally and the timing between transcriptomic dysregulation and behavioral onset. Our wildtype studies saw no overt effects from ∼80% MECP2 mRNA silencing for four months, though early behavioral changes may have been missed. Addressing these questions is likely essential for clinical use. Additionally, plasma biomarkers for target engagement need to be established, with MDS biomarker research from ongoing Ionis Pharmaceuticals trials (ClinicalTrials.gov ID NCT06014541) potentially informing future programs.

This study has developed and preliminarily characterized siRNA compounds for MDS, paving the way for formal preclinical development. These siRNA tools also offer valuable resources for studying MECP2 isoform function and regulation, supporting sustained gene-silencing Rett syndrome models across species. Future work will entail detailed molecular analyses of transcriptomic effects from total and E1 isoform-selective suppression and dose-ranging studies to clarify the pharmacokinetics and pharmacodynamics for safely regulating MECP2 levels.

## MATERIALS AND METHODS

### Study Design

The goal of the study was to identify lead siRNA sequences for isoform E1 selective and total *MECP2* mRNA silencing and to identify optimal chemical scaffolds for *in vivo* potency, durability and tunability. Using standard siRNA design algorithms, we designed a panel of siRNAs targeting all isoforms or isoform E1 of *MECP2.* We selected the size of the panel based on historical hit rates of our in-house algorithm, followed by synthesis and *in vitro* characterization of activity in human HeLa cells as well as mouse N2A cells to identify lead compounds. We further validated target engagement of the lead compounds in both wildtype and humanized MECP2 transgenic mouse models by measuring target mRNA in multiple brain regions at a single dose and two time points. These experiments were done in accordance with approved IACUC protocols at the University of Massachusetts Chan Medical School. The power of the studies was decided based upon historical experience with divalent siRNAs having similar potency in passive uptake experiments *in vitro* which allowed us to estimate effect size and variability for the purposes of power calculations. Once we validated the target engagement of lead siRNAs, we utilized JAX In Vivo Pharmacology Services for behavioral characterization of the Tg1 and Tg3 mouse models, followed by evaluation of behavioral modification of lead compounds following single dose administration at six weeks of age. We selected the dose based on previously published data on ASOs for the treatment of MDS as a benchmark. A single dose rather than multi-dose design was selected based on the observed duration of effect in wildtype mice and the expectation that the durability of a single dose was sufficient to study the effect of the test articles on disease progression in these mice. We opted to measure behavioral endpoints and survival of the Tg1 and Tg3 cohorts rather than schedule euthanasia and perform molecular biology analysis because we already demonstrated target engagement in wildtype mice and did not expect target engagement to be any different in the Tg1 and Tg3 mice. We further decided that behavior and mortality were more relevant to clinical translation than a molecular biology analysis of gene and protein levels.

All graphs were created and analyzed statistically using GraphPad Prism software.

### siRNA Selection

siRNA sequences were selected based on established rules of thermodynamics, G+C content, self-complementarity, and off-targets with seed region homology (*38*). Potential siRNA sequences were further filtered based on isoform selectivity, seed region matches to off-target mRNA and lncRNA and species cross homology. Oligonucleotide synthesis, deprotection, purification for *in vitro* and *in vivo* experiments and LC-MS analysis are included in the Supplementary Materials.

### Cell culture

HeLa cells (ATCC, #CCL-2) and N2A cells (ATCC, #CCL-131) were maintained in Dulbecco′s Modified Eagle′s Medium (DMEM) (Cellgro, #10-013CV). The media were supplemented with 9% fetal bovine serum (FBS) (Gibco, #26140), and cells were grown at 37 °C and 5% CO2. Cells were split every 3 to 7 d and discarded after 15 passages.

### Animal Studies

All experimental studies involving mice were approved by the University of Massachusetts Chan Medical School Institutional Animal Care and Use Committee (IACUC) (protocol # A-2411) and the JAX In Vivo Pharmacology Services IACUC (protocol AUS #21053). Tg1 (JR#8679) and Tg3 (JR#8680) mice were obtained from the Jackson laboratory. Mice were housed at up to five mice per cage (University of Massachusetts Chan Medical School) or 3 mice per cage (JAX In Vivo Pharmacology Services) with food and water ad libitum.

Detailed procedures including surgical administration of test articles and behavioral tests are included in the Supplementary Materials.

### mRNA Quantification

mRNA quantification was performed from *in vitro* cell lysate or from tissue punches stabilized in RNAlater (Invitrogen) using the QuantiGene™ Singleplex branched DNA assay (Invitrogen) as previously described (*57*). Briefly, cells were lysed in QuantiGene™ Lysis Mixture and tissues biopsy punches were homogenized in QuantiGene™ Homogenizing Solution containing 0.2mg/mL Proteinase K according to the manufacturer’s instructions. mRNA was detected according to the QuantiGene™ Singleplex branched DNA assay protocol using the following probe sets: mouse HPRT (SB-15463), mouse MECP2 (SB-11697), human HPRT (SA-10030), human MECP2 (SA-14665). Custom probe sets were designed to selectively detect mouse MECP2 E1 (DR7DPC3), mouse MECP2 E2 (DR47VRG), human MECP2 E1 (DR32Z69) and human MECP2 E2 (DR2W7MC). Luminescence was detected on a Tecan M1000 (Tecan) plate reader.

### Microscopy and immunofluorescence of tissue sections

Formalin-fixed, paraffin-embedded tissue sections were deparaffinized in xylene, rehydrated, and stained with MECP2 (D43) XP Rabbit mAb #3456 at a 1:50 dilution (Cell Signaling Technology). Donkey anti-rabbit IgG (H+L) (Alexa Fluor® 594) (#A-21207, Invitrogen) was used as secondary antibody at a 1:500 dilution. The nucleus was stained with DAPI (Molecular Probes). Images of three sections per mouse, 100 µm apart, were acquired with Leica DMi8 inverted tiling microscope (Leica Microsystems) and processed using LAS X.

### Image analysis and quantification of nuclei

Image analysis of immunofluorescence tissue sections were performed using the ImageJ tools. Briefly, images were loaded onto ImageJ and split into a DAPI channel and a MECP2 channel. The threshold for the DAPI signal was adjusted using the ‘adjust → threshold’ options until the DAPI signal was accurately partitioned from background. After threshold adjustment, the particle analysis function was used by going to ‘analyze → analyze particles’ to obtain a region of interest (ROI) list. Each ROI was a nucleus identified by the ImageJ software. The ROI list was saved as ‘total nuclei’. Next, the MECP2 channel was preprocessed by background subtraction using a rolling ball radius of 50 pixels.

A custom ImageJ script was run to measure the MECP2 signal per nucleus that was previously identified from the DAPI channel and is available in supplementary methods.

## List of Supplementary Materials

Materials and Methods

Fig S1 to S3

Table S1

## Funding

This project was funded by a Rett Syndrome Research Trust grant to AK and National Institutes of Health (NIH) grants R01 HD086111 and S10 OD020012 to AK.

## Author contributions

Conceptualization: VNH, MC, AK

Methodology: VNH, AS, JC, SRH, ZK, DE, CF, AK

Investigation: VNH, AS, JC, DO, QT, DE, NM, DC, JS

Behavioral experiments: AEC-A, LB

Visualization: VNH

Funding acquisition: AK

Project administration: VNH

Supervision: VNH, AK

Writing – original draft: VNH, AK

Writing – review & editing: VNH, AK

## Competing interests

AK and VNH have filed patent applications related to this work. AK discloses ownership of stocks in Advirna and RXi Pharmaceuticals and is a founder of Atalanta Therapeutics and Comanche Biopharma. VNH is an employee of Comanche Biopharma and owns stock options. The remaining authors declare no competing interests.

## Data and materials availability

All data are available in the main text or the supplementary materials.

## Notes

### Competing Interest Statement

The authors have declared no competing interest.

