## Supplementary Information for "Single-dose administration of therapeutic divalent siRNA targeting MECP2 prevents lethality for one year in an MECP2 duplication mouse model"

**Supplementary Materials and Methods**
**Oligonucleotide synthesis**

Oligonucleotides were synthesized by phosphoramidite solid-phase synthesis on automated synthesizer using a MerMade12 (Biosearch Technologies, Novato, CA), Dr Oligo 48 (Biolytic, Fremont, CA) or AKTA Oligopilot 100 (Cytiva, Marlborough, MA). 5'-(E)-Vinyl tetraphosphonate (pivaloyloxymethyl) 2'-O-methyl-uridine 3'-CE phosphoramidite was used for the addition of 5’-Vinyl Phosphonate, 2′-F, 2′-OMe phosphoramidites with standard protecting groups were used to make the modified oligonucleotides. Phosphoramidites were dissolved at 0.1 M in anhydrous acetonitrile (ACN), with added anhydrous 15% dimethylformamide in the case of the 2'-OMe-Uridine amidite. 5-(Benzylthio)-1H-tetrazole (BTT) was used as the activator at 0.25 M. Coupling times were 4 minutes. Detritylations was performed using 3% trichloroacetic acid in dichloromethane or Toluene. Capping reagents used were CAP A (20% N-methylimidazole in ACN) and CAP B (20% acetic anhydride and 30% 2,6-lutidine in ACN). Phosphite oxidation to convert to phosphate or phosphorothioate was performed with 0.05 M iodine in pyridine-water (9:1, v/v) or 0.1 M solution of 3-[(dimethylaminomethylene)amino]-3H-1,2,4-dithiazole-5-thione (DDTT) in pyridine for 3 min, respectively. All reagents were purchased from Chemgenes, Wilmington, MA, phospohoramidites were purchased from Chemgenes and Hongene Biotech, Union City, CA. Oligonucleotides were grown on long-chain alkyl amine (LCAA) 500Å controlled pore glass (CPG) functionalized with Unylinker terminus for unconjugated oligonucleotides. Cholesterol conjugated oligonucleotides were synthesized on a 500Å LCAA-CPG support, tetraethyleneglycol cholesterol moiety was the functionalized group joined by a succinate linker to the support (Chemgenes). Divalent oligonucleotides were synthesized on a 1000Å LCAA-CPG support functionalized via succinyl linker with a glycerol-tetraethyleneglycol linker (Hongene Biotech).

**Deprotection and purification of oligonucleotides for in vitro experiments**

Oligonucleotides with or without cholesterol conjugate were cleaved and deprotected on-column with Ammonia gas (Airgas Specialty Gases). Briefly, columns were pre-wet with 100uL of water and immediately spun to remove the excess water. Columns were then placed in a reaction chamber (Biolytic) 90min at 65°C. A modified on-column ethanol precipitation protocol was used for desalting and counterion exchange. Briefly, 1mL of 0.1M sodium acetate in 80% ethanol is flushed through the column, followed by a rinse with 1mL 80% ethanol and finally after drying the excess ethanol, oligonucleotides were eluted with 600uL of water in 96 deep well plates.

**Deprotection and Purification of Oligonucleotides for in vivo Experiments**

5’-(E)-Vinyl-phosphonate containing oligonucleotides were cleaved and deprotected with 3% diethylamine in ammonium hydroxide, for 20 h at 35°C with agitation. Divalent oligonucleotides were cleaved and deprotected with 1:1 ammonium hydroxide and 40% aqueous monomethylamine, for 2 h at 25°C with slight agitation. The controlled pore glass was subsequently filtered and rinsed with 30 mL of 5% ACN in water and dried overnight by centrifugal vacuum concentration. Purifications were performed on an Agilent 1290 Infinity II HPLC system using Source 15Q anion exchange resin (Cytiva). The loading solution was 20 mM sodium acetate in 10% ACN in water, and elution solution was the loading solution with 1M sodium bromide. Both oligonucleotide strands were eluted using a linear gradient from 30 to 70% in 40 min at 50°C. Peaks were monitored at 260nm. Pure fractions were combined and desalted by size exclusion using Sephadex G-25 resins (Cytiva). Oligonucleotides were then lyophilized and resuspended in water.

**LC-MS analysis of oligonucleotides**

Purity and identity of oligonucleotides were confirmed by IP-RP HPLC coupled to an Agilent 6530 Accurate-mass Q-TOF. LC parameters: buffer A: 100 mM 1,1,1,3,3,3-hexafluoroisopropanol (HFIP) (Oakwood Chemicals) and 9 mM triethylamine (TEA) (Fisher Scientific) in LC–MS grade water (Fisher Scientific); buffer B:100 mM HFIP and 9 mM TEA in LC–MS grade methanol (Fisher Scientific); column, Agilent AdvanceBio oligonucleotides C18; linear gradient 5–35% B 5min was used for unconjugated and divalent oligonucleotides; linear gradient 25–80% B 5min was used for cholesterol conjugated oligonucleotides; temperature, 60°C; flow rate, 0.85 ml/min. Peaks were monitored at 260nm. MS parameters: Source, electrospray ionization; ion polarity, negative mode; range, 100–3,200 m/z; scan rate, 2 spectra/s; capillary voltage, 4,000; fragmentor, 200 V; gas temp, 325°C.

**ImageJ script for quantification of MECP2 in nuclei**

```
//select directory to save output

 dir = getDirectory("Choose a Directory");
waitForUser("click on image you wish to analyze. make sure you have ROI set of nuclei already loaded. click ok to continue");

//calling ROI list of nuclei
n = roiManager('count');
for (i = 0; i < n; i++) {
roiManager('select', i);

//process roi here

run("Set Measurements...", "area mean modal min integrated median redirect=None decimal=3");
run("Measure");

//selectWindow("Summary");
}
selectWindow("Results");
Table.save(dir+File.separator+"results"+i+".csv"); //saves table
 waitForUser ("done, close Results table before next analysis.");
```

**Animal Studies at JAX In Vivo Pharmacology Services**

**In-life Observations and Procedures**

**Complete Blinding of the study**

Cage Cards, test article containers and Experimental Logs were blinded for treatment as “A” (Test article NTC), “B” (Test article M2), “C” (Test article M3), to blind the technicians performing injections, phenotypic scoring, body weight recording and behavioral tests for the dose injected. In addition, genotypes were also blinded.

**Intracerebral Ventricular Injections (ICV) in Adult Mice**

The mouse was anesthetized, and analgesia is administered.

The fur was removed, using clippers from the dorsal head in a roughly triangular area beginning ~ 3mm caudal to the nares and extending caudally to the cervical vertebrae. The lateral borders follow a line from the eye to the base of the pinna. Loose fur was removed with adhesive tape, dry gauze or gauze slightly dampened with ethanol. Ophthalmic ointment was placed on the eyes to prevent drying of the cornea. The skin was disinfected with a surgical scrub (iodine or chlorhexidine) and 70% ethanol using sterile swabs. Application of 70% ethanol starts in the center of the proposed incision site and works outward in ever widening circles to cover the entire clipped area. Using a new sterile swab, the surgical scrub was applied in the same manner. Repeat 70% ethanol and surgical scrub one additional time. The mouse was placed in ventral recumbency with the head secured on a stereotaxic device. A 1cm midsagittal skin incision was made on the scalp to expose the skull. The underlying fascia was separated, and the skin was retracted laterally. With a sterile cotton-tipped applicator soaked in 3% hydrogen peroxide, the skull was cleared of tissue down to the bone. The site was flushed with saline. The sterile drill was attached to the manipulator arm of the stereotaxic unit and the tip of the drill bit was aligned directly over bregma, a landmark on the skull and a stereotaxic reference point. Bregma was the position on the skull where the coronal and sagittal sutures intersect. The surgeon sets the X and Y coordinates of the stereotaxic device to zero. All subsequent stereotaxic coordinates reference these zero points. The manipulator arm with the attached drill is positioned at M/L +/-1.0, A/P -.40, and depth of 2.6 for injection (depth varies from 2.4 to 2.8). A hole was drilled through the skull but leaving the dura intact. The diameter of the drill bur was generally 0.6-1.6 mm. The site is flushed with sterile saline. The drill was removed and replaced on the manipulator arm with the sterile injector. The injector was lowered through the hole in the skull to the predetermined Z point. The siRNA was slowly injected, and the injector held in place for ~1 minute. Mice will receive a bilateral injection of 5 ul per ventricle for a total of 10 ul per mouse. 0.1% bupivacaine was applied to edges of the skin incision. The skin was closed with 6-0 suture with a swaged-on needle or tissue adhesive. Absorbable suture may be used to close skin incisions, but it was not preferred as it causes a greater inflammatory response than non-absorbable suture. Silk, plain gut, or chromic gut suture must never be used to close skin incisions. All wound closure material, except tissue adhesive or absorbable suture, must be removed after the wound has healed.

**Open Field Test**

Subjects were acclimated to the testing room for at least 60 min.

Equipment: Omnitech Versamax Open Field Arenas (40cmx 40cm x 40cm) with photobeams.

Mice were placed individually into the center of the arena. Data were recorded via a sensitive infrared (IR) photobeam three-dimensional grid system that is invisible to mice. When the mouse moves or travels, its body breaks the otherwise continuous beam. The automated system then translates the beam breaks into measurements such as distance traveled (cm), horizontal activity, vertical activity (rearing), and time spent in the center (anxiety-like) for 60 min.

**Rotarod**

Subjects were acclimated to the testing room for a minimum of 60 min.

Equipment: Ugo-Basile Accelerating Rotarod for Mice (model 47600).

Up to 5 mice were placed on the rod, which was rotating at the baseline speed of 5 RPM. Once all mice were placed on the rod, it accelerated linearly to 40 RPM over a period of 300 sec (5 min). Mice are run on three successive trials, and the latency to fall, which is measured as the time from the initiation of the 300 sec linear acceleration period to when the mouse falls off the rod into the tray below, is recorded for each mouse on each trial. Mice lasting 300 sec on the rod, without falling, receive a “latency to fall” time of 300 sec.

**Nest building**

Nesting behavior is an intrinsic behavior performed by male and female rodents that requires fine motor skills. The nest building test is a simple and versatile behavioral test suitable for the evaluation of motor deficits, cognitive decline as well as changes in general health or welfare. It is sensitive to environmental and physiological challenges, genetic mutations, and pharmacological interventions. There are several advantages of this test: it is performed in the home cage, without the presence of the experimenter, bedding material that is not used can be weighed to obtain an objective measure, and it is simple to perform while still yielding robust readouts. To test the individual nest building behavior, mice were individually housed in cages containing wood chip bedding and one square of pressed cotton `nestlet´. No other nesting material was provided. After 24hrs, photographs of the nests were taken, un-manipulated bedding material was weighed, and the manipulation of the nestlet and the quality of the built nest were analyzed manually according to a five-point scale.

**Supplementary Figures**

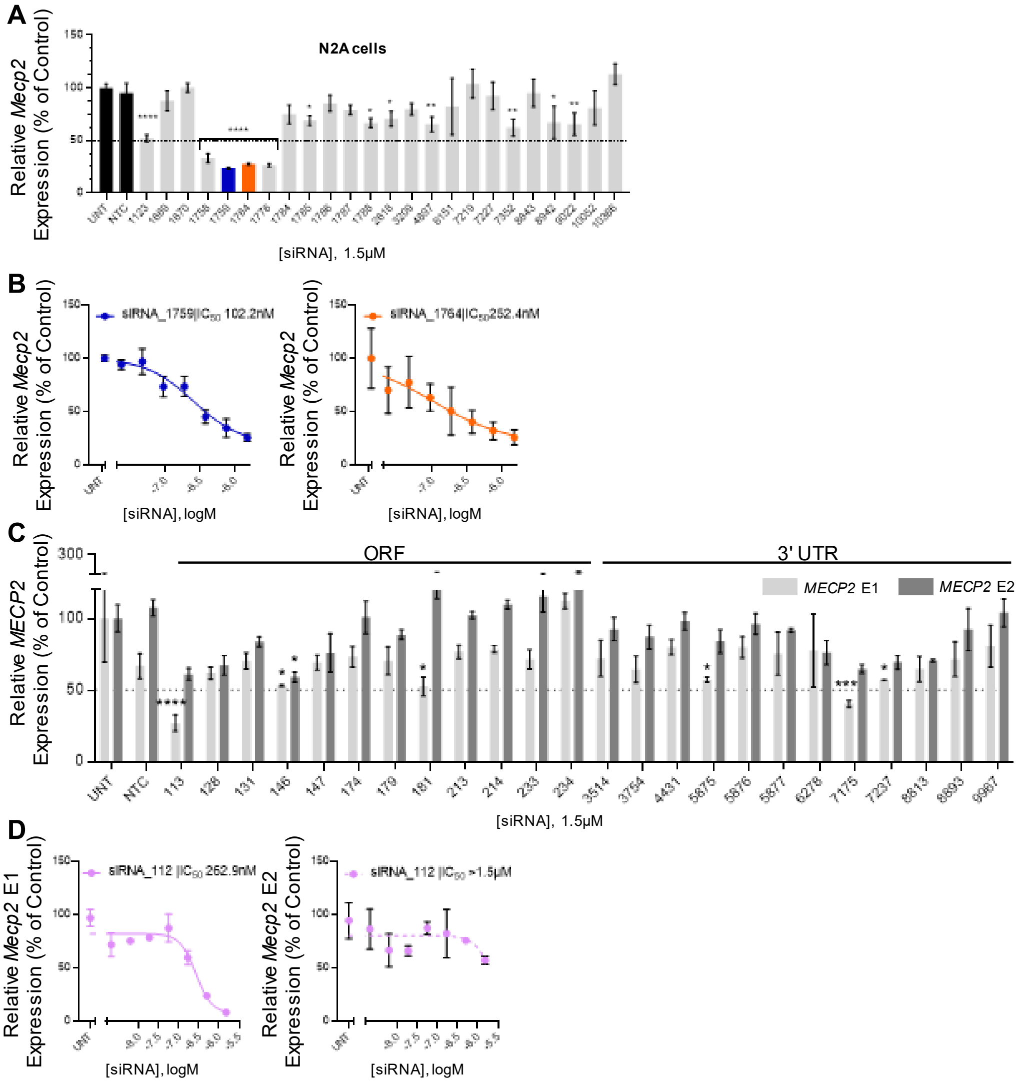

**Supplementary Figure S1**. **Lead siRNAs identified through in vitro screening**. (A) Relative *Mecp2* mRNA expression in mouse Neuro2A cells 72h post treatment with siRNAs at 1.5µM. (B) Relative *Mecp2* mRNA expression in mouse Neuro2A cells 72h post treatment with lead siRNAs in a seven-point dose response. (C) Relative *Mecp2* E1 (solid bars) and *Mecp2* E2 (patterned bars) mRNA expression in mouse Neuro2A cells 72h post treatment with siRNAs at 1.5µM. (D) Relative *Mecp2* E1 (left) and E2 (right) mRNA expression in mouse Neuro2A cells 72h post treatment with siRNA_112 in a seven-point dose response. Gene expression was measured using the QuantiGene™ Singleplex Assay Kit. Data represented as mean±s.e.m. of three independent replicates. Statistical analysis performed using Ordinary one-way ANOVA with Dunnett’s adjustment for multiple comparisons.

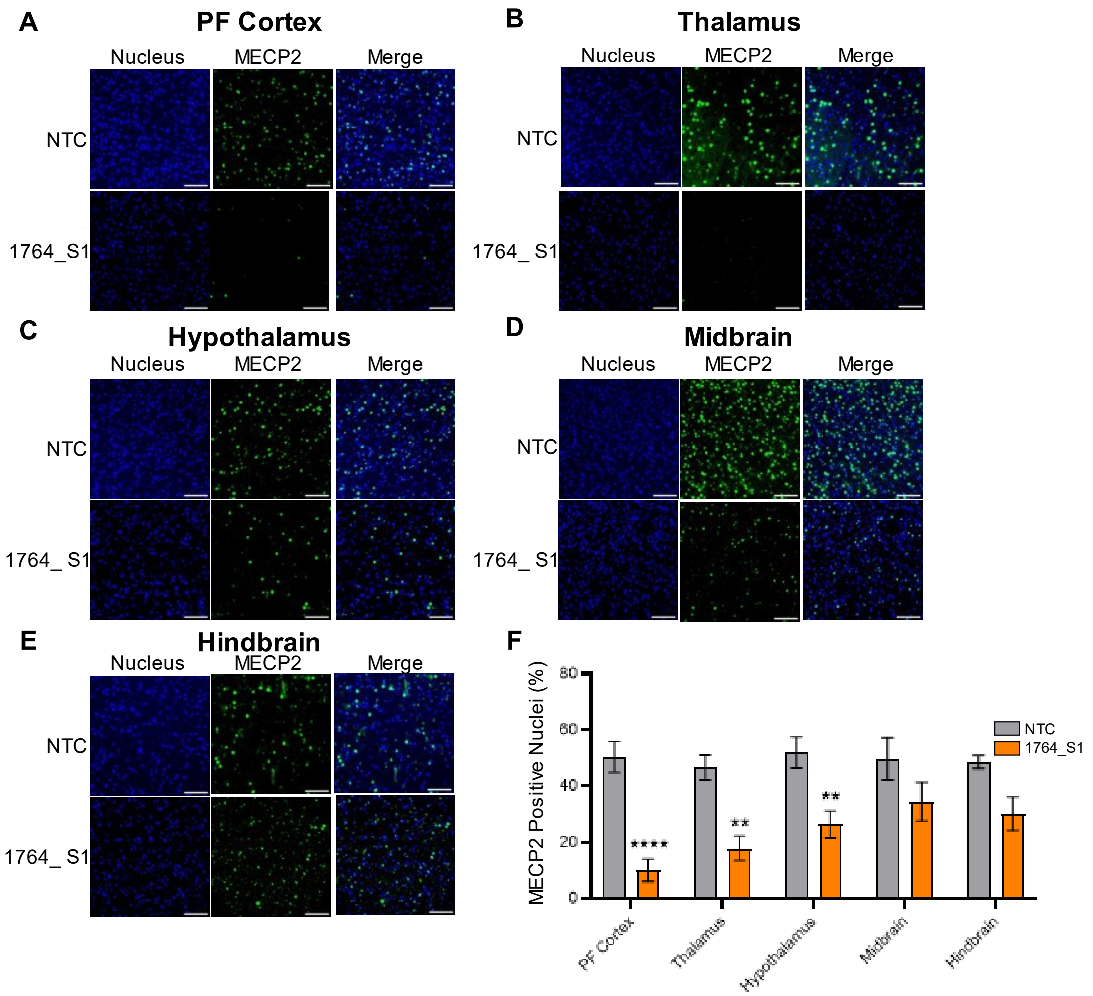

**Supplementary Figure S2. Quantification of MECP2 positive nuclei in brain sections.** Representative immunofluorescence images of DAPI and MECP2 stained sections of (A) prefrontal (PF) cortex, (B) thalamus, (C) hypothalamus, (D) midbrain, and (E) hindbrain along with (F) quantification of MECP2 positive nuclei. Data represented as mean±s.e.m. of individual animals. Immunofluorescence imaging was performed on sagittal brain sections stained with DAPI and anti-MECP2 antibody and imaged using Leica DMi8 widefield microscope. Quantification of MECP2 positive nuclei was performed using a custom imageJ script. Statistical analysis performed using two-way ANOVA with Dunnett’s adjustment for multiple comparisons (**-p<0.01, ****-p<0.0001).

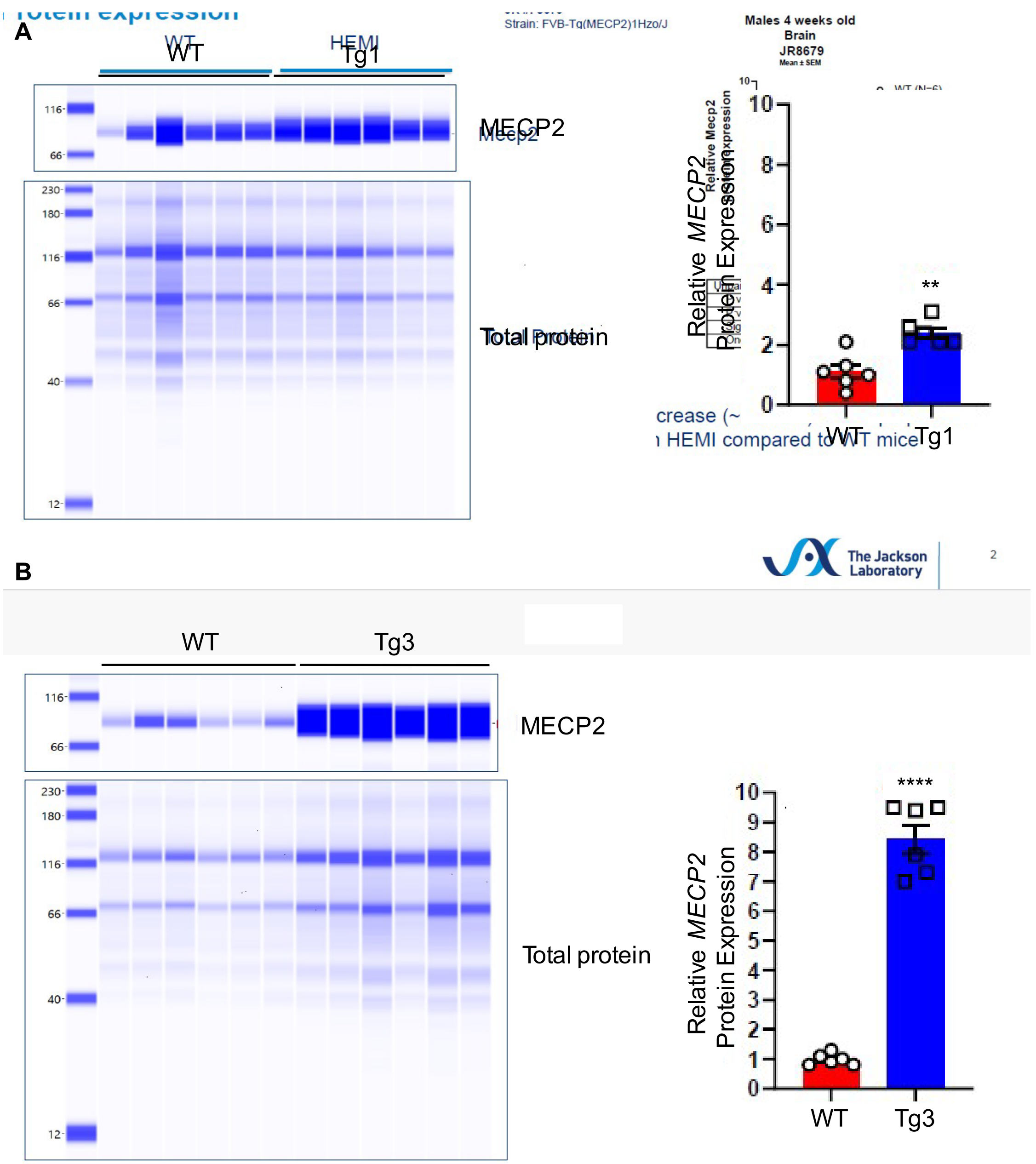

**Supplementary Figure S3. Validation of Tg1 and Tg3 mouse models.** Western blot gel images (left) and quantification of protein bands (right) from 4 weeks old (A) Tg1 mice and (B) Tg3 mice. Mecp2 protein bands were normalized to total protein. WT = FVB background, HEMI = transgenic strain. Data in bar graphs represented as mean±s.d. of individual animals (n=6). Statistical significance computed by unpaired t-test (**-p<0.01, ****-p<0.0001).

**Supplementary Table S1. List of oligonucleotide sequences used in this study**

| Single strand ID | Oligo Name | Modified sequence |
|---|---|---|
| 7051 | MECP2_575_s | (mG)#(mA)#(mA)(mG)(fU)(fA)(fU)(mG)(fA)(mU)(mG)(mU)(mG)#(mU)#(mA)-TegChol |
| 7087 | MECP2_575_as | P(mU)#(fA)#(mC)(mA)(mC)(fA)(mU)(mC)(mA)(mU)(mA)(mC)(mU)#(fU)#(mC)#(fC)#(mC)#(mA)#(mG)#(fC) |
| 7052 | MECP2_572_s | (mU)#(mG)#(mG)(mG)(fA)(fA)(fG)(mU)(fA)(mU)(mG)(mA)(mU)#(mG)#(mA)-TegChol |
| 7088 | MECP2_572_as | P(mU)#(fC)#(mA)(mU)(mC)(fA)(mU)(mA)(mC)(mU)(mU)(mC)(mC)#(fC)#(mA)#(fG)#(mC)#(mA)#(mG)#(fA) |
| 7053 | MECP2_300_s | (mA)#(mA)#(mG)(mU)(fU)(fU)(fA)(mA)(fA)(mA)(mA)(mG)(mG)#(mU)#(mA)-TegChol |
| 7089 | MECP2_300_as | P(mU)#(fA)#(mC)(mC)(mU)(fU)(mU)(mU)(mU)(mA)(mA)(mA)(mC)#(fU)#(mU)#(fG)#(mA)#(mG)#(mG)#(fG) |
| 7054 | MECP2_827_s | (mC)#(mA)#(mC)(mG)(fU)(fC)(fA)(mG)(fA)(mG)(mG)(mG)(mU)#(mG)#(mA)-TegChol |
| 7090 | MECP2_827_as | P(mU)#(fC)#(mA)(mC)(mC)(fC)(mU)(mC)(mU)(mG)(mA)(mC)(mG)#(fU)#(mG)#(fG)#(mC)#(mC)#(mG)#(fC) |
| 7055 | MECP2_603_s | (mC)#(mA)#(mG)(mG)(fG)(fA)(fA)(mA)(fA)(mG)(mC)(mC)(mU)#(mU)#(mA)-TegChol |
| 7091 | MECP2_603_as | P(mU)#(fA)#(mA)(mG)(mG)(fC)(mU)(mU)(mU)(mU)(mC)(mC)(mC)#(fU)#(mG)#(fG)#(mG)#(mG)#(mA)#(fU) |
| 7056 | MECP2_887_s | (mC)#(mA)#(mA)(mG)(fA)(fU)(fG)(mC)(fC)(mU)(mU)(mU)(mU)#(mC)#(mA)-TegChol |
| 7092 | MECP2_887_as | P(mU)#(fG)#(mA)(mA)(mA)(fA)(mG)(mG)(mC)(mA)(mU)(mC)(mU)#(fU)#(mG)#(fA)#(mC)#(mA)#(mA)#(fG) |
| 7057 | MECP2_285_s | (mA)#(mA)#(mG)(mG)(fA)(fC)(fA)(mA)(fA)(mC)(mC)(mC)(mC)#(mU)#(mA)-TegChol |
| 7093 | MECP2_285_as | P(mU)#(fA)#(mG)(mG)(mG)(fG)(mU)(mU)(mU)(mG)(mU)(mC)(mC)#(fU)#(mU)#(fG)#(mA)#(mG)#(mG)#(fC) |
| 7058 | MECP2_605_s | (mG)#(mG)#(mG)(mA)(fA)(fA)(fA)(mG)(fC)(mC)(mU)(mU)(mU)#(mC)#(mA)-TegChol |
| 7094 | MECP2_605_as | P(mU)#(fG)#(mA)(mA)(mA)(fG)(mG)(mC)(mU)(mU)(mU)(mU)(mC)#(fC)#(mC)#(fU)#(mG)#(mG)#(mG)#(fG) |
| 7059 | MECP2_662_s | (mC)#(mA)#(mC)(mA)(fU)(fC)(fC)(mC)(fU)(mG)(mG)(mA)(mC)#(mC)#(mA)-TegChol |

| | | |
|---|---|---|
| 7095 | MECP2_662_as | P(mU)#(fG)#(mG)(mU)(mC)(fC)(mA)(mG)(mG)(mG)(mA)(mU)(mG)#(fU)#(mG)#(fU)#(mC)#(mG)#(mC)#(fC) |
| 7060 | MECP2_535_s | (mC)#(mA)#(mC)(mG)(fG)(fA)(fA)(mG)(fC)(mU)(mU)(mA)(mA)#(mG)#(mA)-TegChol |
| 7096 | MECP2_535_as | P(mU)#(fC)#(mU)(mU)(mA)(fA)(mG)(mC)(mU)(mU)(mC)(mC)(mG)#(fU)#(mG)#(fU)#(mC)#(mC)#(mA)#(fG) |
| 7061 | MECP2_303_s | (mU)#(mU)#(mU)(mA)(fA)(fA)(fA)(mA)(fG)(mG)(mU)(mG)(mA)#(mA)#(mA)-TegChol |
| 7097 | MECP2_303_as | P(mU)#(fU)#(mU)(mC)(mA)(fC)(mC)(mU)(mU)(mU)(mU)(mU)(mA)#(fA)#(mA)#(fC)#(mU)#(mU)#(mG)#(fA) |
| 7062 | MECP2_895_s | (mC)#(mU)#(mU)(mU)(fU)(fC)(fA)(mA)(fA)(mC)(mU)(mU)(mC)#(mG)#(mA)-TegChol |
| 7098 | MECP2_895_as | P(mU)#(fC)#(mG)(mA)(mA)(fG)(mU)(mU)(mU)(mG)(mA)(mA)(mA)#(fA)#(mG)#(fG)#(mC)#(mA)#(mU)#(fC) |
| 7063 | MECP2_1776_s | (mA)#(mU)#(mU)(mA)(fA)(fC)(fU)(mG)(fA)(mA)(mA)(mU)(mA)#(mA)#(mA)-TegChol |
| 7099 | MECP2_1776_as | P(mU)#(fU)#(mU)(mA)(mU)(fU)(mU)(mC)(mA)(mG)(mU)(mU)(mA)#(fA)#(mU)#(fC)#(mG)#(mG)#(mG)#(fA) |
| 7064 | MECP2_1785_s | (mA)#(mA)#(mU)(mA)(fA)(fA)(fA)(mA)(fA)(mU)(mA)(mU)(mU)#(mU)#(mA)-TegChol |
| 7100 | MECP2_1785_as | P(mU)#(fA)#(mA)(mA)(mU)(fA)(mU)(mU)(mU)(mU)(mU)(mU)(mA)#(fU)#(mU)#(fU)#(mC)#(mA)#(mG)#(fU) |
| 7065 | MECP2_1759_s | (mU)#(mC)#(mU)(mG)(fA)(fC)(fA)(mA)(fA)(mG)(mC)(mU)(mU)#(mC)#(mA)-TegChol |
| 7101 | MECP2_1759_as | P(mU)#(fG)#(mA)(mA)(mG)(fC)(mU)(mU)(mU)(mG)(mU)(mC)(mA)#(fG)#(mA)#(fG)#(mC)#(mC)#(mC)#(fU) |
| 7066 | Mecp2_1123_s | (mC)#(mA)#(mA)(mA)(fA)(fA)(fG)(mA)(fA)(mA)(mG)(mC)(mC)#(mG)#(mA)-TegChol |
| 7102 | Mecp2_1123_as | P(mU)#(fC)#(mG)(mG)(mC)(fU)(mU)(mU)(mC)(mU)(mU)(mU)(mU)#(fU)#(mG)#(fG)#(mC)#(mC)#(mU)#(fC) |
| 7067 | MECP2_1787_s | (mU)#(mA)#(mA)(mA)(fA)(fA)(fA)(mU)(fA)(mU)(mU)(mU)(mU)#(mU)#(mA)-TegChol |
| 7103 | MECP2_1787_as | P(mU)#(fA)#(mA)(mA)(mA)(fA)(mU)(mA)(mU)(mU)(mU)(mU)(mU)#(fU)#(mA)#(fU)#(mU)#(mU)#(mC)#(fA) |
| 7068 | MECP2_1784_s | (mA)#(mA)#(mA)(mU)(fA)(fA)(fA)(mA)(fA)(mA)(mU)(mA)(mU)#(mU)#(mA)-TegChol |
| 7104 | MECP2_1784_as | P(mU)#(fA)#(mA)(mU)(mA)(fU)(mU)(mU)(mU)(mU)(mU)(mA)(mU)#(fU)#(mU)#(fC)#(mA)#(mG)#(mU)#(fU) |

| | | |
|---|---|---|
| 7069 | MECP2_1786_s | (mA)#(mU)#(mA)(mA)(fA)(fA)(fA)(mA)(fU)(mA)(mU)(mU)(mU)#(mU)#(mA)-TegChol |
| 7105 | MECP2_1786_as | P(mU)#(fA)#(mA)(mA)(mA)(fU)(mA)(mU)(mU)(mU)(mU)(mU)(mU)#(fA)#(mU)#(fU)#(mU)#(mC)#(mA)#(fG) |
| 7070 | MECP2_1788_s | (mA)#(mA)#(mA)(mA)(fA)(fA)(fU)(mA)(fU)(mU)(mU)(mU)(mU)#(mU)#(mA)-TegChol |
| 7106 | MECP2_1788_as | P(mU)#(fA)#(mA)(mA)(mA)(fA)(mA)(mU)(mA)(mU)(mU)(mU)(mU)#(fU)#(mU)#(fA)#(mU)#(mU)#(mU)#(fC) |
| 7071 | MECP2_1758_s | (mC)#(mU)#(mC)(mU)(fG)(fA)(fC)(mA)(fA)(mA)(mG)(mC)(mU)#(mU)#(mA)-TegChol |
| 7107 | MECP2_1758_as | P(mU)#(fA)#(mA)(mG)(mC)(fU)(mU)(mU)(mG)(mU)(mC)(mA)(mG)#(fA)#(mG)#(fC)#(mC)#(mC)#(mU)#(fA) |
| 7072 | MECP2_1669_s | (mA)#(mU)#(mA)(mU)(fU)(fU)(fU)(mU)(fU)(mU)(mU)(mU)(mC)#(mU)#(mA)-TegChol |
| 7108 | MECP2_1669_as | P(mU)#(fA)#(mG)(mA)(mA)(fA)(mA)(mA)(mA)(mA)(mA)(mA)(mU)#(fA)#(mU)#(fU)#(mU)#(mU)#(mU)#(fU) |
| 7073 | MECP2_1670_s | (mU)#(mA)#(mU)(mU)(fU)(fU)(fU)(mU)(fU)(mU)(mU)(mC)(mU)#(mU)#(mA)-TegChol |
| 7109 | MECP2_1670_as | P(mU)#(fA)#(mA)(mG)(mA)(fA)(mA)(mA)(mA)(mA)(mA)(mA)(mA)#(fU)#(mA)#(fU)#(mU)#(mU)#(mU)#(fU) |
| 7074 | MECP2_1764_s | (mC)#(mA)#(mA)(mA)(fG)(fC)(fU)(mU)(fC)(mC)(mC)(mG)(mA)#(mU)#(mA)-TegChol |
| 7110 | MECP2_1764_as | P(mU)#(fA)#(mU)(mC)(mG)(fG)(mG)(mA)(mA)(mG)(mC)(mU)(mU)#(fU)#(mG)#(fU)#(mC)#(mA)#(mG)#(fA) |
| 7075 | Mecp2_4897_s | (mC)#(mA)#(mU)(mU)(fG)(fU)(fA)(mA)(fA)(mG)(mU)(mG)(mU)#(mG)#(mA)-TegChol |
| 7111 | Mecp2_4897_as | P(mU)#(fC)#(mA)(mC)(mA)(fC)(mU)(mU)(mU)(mA)(mC)(mA)(mA)#(fU)#(mG)#(fU)#(mU)#(mC)#(mA)#(fA) |
| 7076 | Mecp2_7219_s | (mU)#(mA)#(mU)(mA)(fU)(fA)(fU)(mA)(fU)(mA)(mU)(mC)(mU)#(mG)#(mA)-TegChol |
| 7112 | Mecp2_7219_as | P(mU)#(fC)#(mA)(mG)(mA)(fU)(mA)(mU)(mA)(mU)(mA)(mU)(mA)#(fU)#(mA)#(fU)#(mA)#(mU)#(mA)#(fU) |
| 7077 | Mecp2_10052_s | (mU)#(mU)#(mU)(mG)(fG)(fG)(fA)(mC)(fA)(mA)(mU)(mU)(mA)#(mC)#(mA)-TegChol |
| 7113 | Mecp2_10052_as | P(mU)#(fG)#(mU)(mA)(mA)(fU)(mU)(mG)(mU)(mC)(mC)(mC)(mA)#(fA)#(mA)#(fG)#(mC)#(mA)#(mC)#(fA) |

| | | |
|---|---|---|
| 7078 | Mecp2_6151_s | (mA)#(mA)#(mA)(mA)(fA)(fU)(fU)(mA)(fU)(mA)(mU)(mU)(mU)#(mU)#(mA)-TegChol |
| 7114 | Mecp2_6151_as | P(mU)#(fA)#(mA)(mA)(mA)(fU)(mA)(mU)(mA)(mA)(mU)(mU)(mU)#(fU)#(mU)#(fU)#(mG)#(mU)#(mU)#(fA) |
| 7079 | Mecp2_3209_s | (mC)#(mC)#(mU)(mU)(fA)(fA)(fA)(mC)(fA)(mA)(mU)(mG)(mA)#(mG)#(mA)-TegChol |
| 7115 | Mecp2_3209_as | P(mU)#(fC)#(mU)(mC)(mA)(fU)(mU)(mG)(mU)(mU)(mU)(mA)(mA)#(fG)#(mG)#(fU)#(mC)#(mU)#(mG)#(fA) |
| 7080 | Mecp2_2618_s | (mA)#(mA)#(mG)(mG)(fA)(fA)(fA)(mA)(fU)(mA)(mU)(mU)(mC)#(mU)#(mA)-TegChol |
| 7116 | Mecp2_2618_as | P(mU)#(fA)#(mG)(mA)(mA)(fU)(mA)(mU)(mU)(mU)(mU)(mC)(mC)#(fU)#(mU)#(fU)#(mC)#(mU)#(mU)#(fC) |
| 7081 | Mecp2_9022_s | (mC)#(mA)#(mG)(mA)(fG)(fA)(fC)(mA)(fA)(mA)(mU)(mG)(mC)#(mU)#(mA)-TegChol |
| 7117 | Mecp2_9022_as | P(mU)#(fA)#(mG)(mC)(mA)(fU)(mU)(mU)(mG)(mU)(mC)(mU)(mC)#(fU)#(mG)#(fG)#(mA)#(mA)#(mC)#(fA) |
| 7082 | Mecp2_7352_s | (mA)#(mA)#(mG)(mG)(fG)(fA)(fA)(mA)(fA)(mA)(mA)(mU)(mU)#(mU)#(mA)-TegChol |
| 7118 | Mecp2_7352_as | P(mU)#(fA)#(mA)(mA)(mU)(fU)(mU)(mU)(mU)(mU)(mC)(mC)(mC)#(fU)#(mU)#(fG)#(mU)#(mC)#(mC)#(fU) |
| 7083 | Mecp2_10366_s | (mC)#(mA)#(mU)(mA)(fA)(fG)(fG)(mU)(fU)(mC)(mU)(mU)(mU)#(mU)#(mA)-TegChol |
| 7119 | Mecp2_10366_as | P(mU)#(fA)#(mA)(mA)(mA)(fG)(mA)(mA)(mC)(mC)(mU)(mU)(mA)#(fU)#(mG)#(fA)#(mA)#(mA)#(mA)#(fA) |
| 7084 | Mecp2_8843_s | (mU)#(mU)#(mG)(mG)(fU)(fU)(fU)(mU)(fA)(mU)(mU)(mU)(mU)#(mU)#(mA)-TegChol |
| 7120 | Mecp2_8843_as | P(mU)#(fA)#(mA)(mA)(mA)(fA)(mU)(mA)(mA)(mA)(mA)(mC)(mC)#(fA)#(mA)#(fA)#(mC)#(mC)#(mA)#(fA) |
| 7085 | Mecp2_8942_s | (mG)#(mU)#(mG)(mA)(fA)(fA)(fG)(mG)(fA)(mA)(mC)(mU)(mU)#(mU)#(mA)-TegChol |
| 7121 | Mecp2_8942_as | P(mU)#(fA)#(mA)(mA)(mG)(fU)(mU)(mC)(mC)(mU)(mU)(mU)(mC)#(fA)#(mC)#(fC)#(mC)#(mA)#(mC)#(fC) |
| 7086 | Mecp2_7227_s | (mU)#(mA)#(mU)(mC)(fU)(fG)(fU)(mA)(fU)(mA)(mU)(mU)(mU)#(mC)#(mA)-TegChol |
| 7122 | Mecp2_7227_as | P(mU)#(fG)#(mA)(mA)(mA)(fU)(mA)(mU)(mA)(mC)(mA)(mG)(mA)#(fU)#(mA)#(fU)#(mA)#(mU)#(mA)#(fU) |
| 8174 | MECP2_E1_s_111 | (mA)#(mG)#(fG)(mA)(fG)(mG)(fA)(mG)(fA)(mG)(mA)(mC)(fU)#(mG)#(mA)-TegChol |

| | | |
|---|---|---|
| 8222 | MECP2_E1_as_111 | P(mU)#(fC)#(mA)(fG)(fU)(fC)(mU)(fC)(mU)(fC)(mC)(fU)(mC)#(fC)#(mU)#(fC)#(mG)#(mC)#(mC)#(fU) |
| 8168 | MECP2_E1_s_112 | (mG)#(mG)#(fA)(mG)(fG)(mA)(fG)(mA)(fG)(mA)(mC)(mU)(fG)#(mG)#(mA)-TegChol |
| 8216 | MECP2_E1_as_112 | P(mU)#(fC)#(mC)(fA)(fG)(fU)(mC)(fU)(mC)(fU)(mC)(fC)(mU)#(fC)#(mC)#(fU)#(mC)#(mG)#(mC)#(fC) |
| 8169 | MECP2_E1_s_113 | (mG)#(mA)#(fG)(mG)(fA)(mG)(fA)(mG)(fA)(mC)(mU)(mG)(fG)#(mA)#(mA)-TegChol |
| 8217 | MECP2_E1_as_113 | P(mU)#(fU)#(mC)(fC)(fA)(fG)(mU)(fC)(mU)(fC)(mU)(fC)(mC)#(fU)#(mC)#(fC)#(mU)#(mC)#(mG)#(fC) |
| 8170 | MECP2_E1_s_114 | (mA)#(mG)#(fG)(mA)(fG)(mA)(fG)(mA)(fC)(mU)(mG)(mG)(fA)#(mA)#(mA)-TegChol |
| 8218 | MECP2_E1_as_114 | P(mU)#(fU)#(mU)(fC)(fC)(fA)(mG)(fU)(mC)(fU)(mC)(fU)(mC)#(fC)#(mU)#(fC)#(mC)#(mU)#(mC)#(fG) |
| 8173 | MECP2_E1_s_115 | (mG)#(mG)#(fA)(mG)(fA)(mG)(fA)(mC)(fU)(mG)(mG)(mA)(fA)#(mG)#(mA)-TegChol |
| 8221 | MECP2_E1_as_115 | P(mU)#(fC)#(mU)(fU)(fC)(fC)(mA)(fG)(mU)(fC)(mU)(fC)(mU)#(fC)#(mC)#(fU)#(mC)#(mC)#(mU)#(fC) |
| 8176 | MECP2_E1_s_117 | (mA)#(mG)#(fA)(mG)(fA)(mC)(fU)(mG)(fG)(mA)(mA)(mG)(fA)#(mA)#(mA)-TegChol |
| 8224 | MECP2_E1_as_117 | P(mU)#(fU)#(mU)(fC)(fU)(fU)(mC)(fC)(mA)(fG)(mU)(fC)(mU)#(fC)#(mU)#(fC)#(mC)#(mU)#(mC)#(fC) |
| 8175 | MECP2_E1_s_118 | (mG)#(mA)#(fG)(mA)(fC)(mU)(fG)(mG)(fA)(mA)(mG)(mA)(fA)#(mA)#(mA)-TegChol |
| 8223 | MECP2_E1_as_118 | P(mU)#(fU)#(mU)(fU)(fC)(fU)(mU)(fC)(mC)(fA)(mG)(fU)(mC)#(fU)#(mC)#(fU)#(mC)#(mC)#(mU)#(fC) |
| 8171 | MECP2_E1_s_121 | (mA)#(mC)#(fU)(mG)(fG)(mA)(fA)(mG)(fA)(mA)(mA)(mA)(fG)#(mU)#(mA)-TegChol |
| 8219 | MECP2_E1_as_121 | P(mU)#(fA)#(mC)(fU)(fU)(fU)(mU)(fC)(mU)(fU)(mC)(fC)(mA)#(fG)#(mU)#(fC)#(mU)#(mC)#(mU)#(fC) |
| 8166 | MECP2_E1_s_122 | (mC)#(mU)#(fG)(mG)(fA)(mA)(fG)(mA)(fA)(mA)(mA)(mG)(fU)#(mC)#(mA)-TegChol |
| 8214 | MECP2_E1_as_122 | P(mU)#(fG)#(mA)(fC)(fU)(fU)(mU)(fU)(mC)(fU)(mU)(fC)(mC)#(fA)#(mG)#(fU)#(mC)#(mU)#(mC)#(fU) |
| 8177 | MECP2_E1_s_123 | (mU)#(mG)#(fG)(mA)(fA)(mG)(fA)(mA)(fA)(mA)(mG)(mU)(fC)#(mA)#(mA)-TegChol |
| 8225 | MECP2_E1_as_123 | P(mU)#(fU)#(mG)(fA)(fC)(fU)(mU)(fU)(mU)(fC)(mU)(fU)(mC)#(fC)#(mA)#(fG)#(mU)#(mC)#(mU)#(fC) |

| | | |
|---|---|---|
| 8164 | MECP2_E1_s_126 | (mA)#(mA)#(fG)(mA)(fA)(mA)(fA)(mG)(fU)(mC)(mA)(mG)(fA)#(mA)#(mA)-TegChol |
| 8212 | MECP2_E1_as_126 | P(mU)#(fU)#(mU)(fC)(fU)(fG)(mA)(fC)(mU)(fU)(mU)(fU)(mC)#(fU)#(mU)#(fC)#(mC)#(mA)#(mG)#(fU) |
| 8172 | MECP2_E1_s_127 | (mA)#(mG)#(fA)(mA)(fA)(mA)(fG)(mU)(fC)(mA)(mG)(mA)(fA)#(mG)#(mA)-TegChol |
| 8220 | MECP2_E1_as_127 | P(mU)#(fC)#(mU)(fU)(fC)(fU)(mG)(fA)(mC)(fU)(mU)(fU)(mU)#(fC)#(mU)#(fU)#(mC)#(mC)#(mA)#(fG) |
| 8155 | MECP2_E1_s_1678 | (mU)#(mU)#(fU)(mC)(fU)(mU)(fU)(mC)(fA)(mG)(mU)(mA)(fA)#(mA)#(mA)-TegChol |
| 8203 | MECP2_E1_as_1678 | P(mU)#(fU)#(mU)(fU)(fA)(fC)(mU)(fG)(mA)(fA)(mA)(fG)(mA)#(fA)#(mA)#(fA)#(mA)#(mA)#(mA)#(fA) |
| 8167 | MECP2_E1_s_1679 | (mU)#(mU)#(fC)(mU)(fU)(mU)(fC)(mA)(fG)(mU)(mA)(mA)(fA)#(mA)#(mA)-TegChol |
| 8215 | MECP2_E1_as_1679 | P(mU)#(fU)#(mU)(fU)(fU)(fA)(mC)(fU)(mG)(fA)(mA)(fA)(mG)#(fA)#(mA)#(fA)#(mA)#(mA)#(mA)#(fA) |
| 8159 | MECP2_E1_s_1680 | (mU)#(mC)#(fU)(mU)(fU)(mC)(fA)(mG)(fU)(mA)(mA)(mA)(fA)#(mA)#(mA)-TegChol |
| 8207 | MECP2_E1_as_1680 | P(mU)#(fU)#(mU)(fU)(fU)(fU)(mA)(fC)(mU)(fG)(mA)(fA)(mA)#(fG)#(mA)#(fA)#(mA)#(mA)#(mA)#(fA) |
| 8162 | MECP2_E1_s_1681 | (mC)#(mU)#(fU)(mU)(fC)(mA)(fG)(mU)(fA)(mA)(mA)(mA)(fA)#(mA)#(mA)-TegChol |
| 8210 | MECP2_E1_as_1681 | P(mU)#(fU)#(mU)(fU)(fU)(fU)(mU)(fA)(mC)(fU)(mG)(fA)(mA)#(fA)#(mG)#(fA)#(mA)#(mA)#(mA)#(fA) |
| 8158 | MECP2_E1_s_1682 | (mU)#(mU)#(fU)(mC)(fA)(mG)(fU)(mA)(fA)(mA)(mA)(mA)(fA)#(mA)#(mA)-TegChol |
| 8206 | MECP2_E1_as_1682 | P(mU)#(fU)#(mU)(fU)(fU)(fU)(mU)(fU)(mA)(fC)(mU)(fG)(mA)#(fA)#(mA)#(fG)#(mA)#(mA)#(mA)#(fA) |
| 8165 | MECP2_E1_s_1683 | (mU)#(mU)#(fC)(mA)(fG)(mU)(fA)(mA)(fA)(mA)(mA)(mA)(fA)#(mA)#(mA)-TegChol |
| 8213 | MECP2_E1_as_1683 | P(mU)#(fU)#(mU)(fU)(fU)(fU)(mU)(fU)(mU)(fA)(mC)(fU)(mG)#(fA)#(mA)#(fA)#(mG)#(mA)#(mA)#(fA) |
| 8163 | MECP2_E1_s_1684 | (mU)#(mC)#(fA)(mG)(fU)(mA)(fA)(mA)(fA)(mA)(mA)(mA)(fA)#(mA)#(mA)-TegChol |
| 8211 | MECP2_E1_as_1684 | P(mU)#(fU)#(mU)(fU)(fU)(fU)(mU)(fU)(mU)(fU)(mA)(fC)(mU)#(fG)#(mA)#(fA)#(mA)#(mG)#(mA)#(fA) |

| | | |
|---|---|---|
| 8157 | MECP2_E1_s_1685 | (mC)#(mA)#(fG)(mU)(fA)(mA)(fA)(mA)(fA)(mA)(mA)(mA)(fA)#(mA)#(mA)-TegChol |
| 8205 | MECP2_E1_as_1685 | P(mU)#(fU)#(mU)(fU)(fU)(fU)(mU)(fU)(mU)(fU)(mU)(fA)(mC)#(fU)#(mG)#(fA)#(mA)#(mA)#(mG)#(fA) |
| 8160 | MECP2_E1_s_1686 | (mA)#(mG)#(fU)(mA)(fA)(mA)(fA)(mA)(fA)(mA)(mA)(mA)(fA)#(mA)#(mA)-TegChol |
| 8208 | MECP2_E1_as_1686 | P(mU)#(fU)#(mU)(fU)(fU)(fU)(mU)(fU)(mU)(fU)(mU)(fU)(mA)#(fC)#(mU)#(fG)#(mA)#(mA)#(mA)#(fG) |
| 8161 | MECP2_E1_s_1687 | (mG)#(mU)#(fA)(mA)(fA)(mA)(fA)(mA)(fA)(mA)(mA)(mA)(fA)#(mA)#(mA)-TegChol |
| 8209 | MECP2_E1_as_1687 | P(mU)#(fU)#(mU)(fU)(fU)(fU)(mU)(fU)(mU)(fU)(mU)(fU)(mU)#(fA)#(mC)#(fU)#(mG)#(mA)#(mA)#(fA) |
| 8156 | MECP2_E1_s_1688 | (mU)#(mA)#(fA)(mA)(fA)(mA)(fA)(mA)(fA)(mA)(mA)(mA)(fA)#(mA)#(mA)-TegChol |
| 8204 | MECP2_E1_as_1688 | P(mU)#(fU)#(mU)(fU)(fU)(fU)(mU)(fU)(mU)(fU)(mU)(fU)(mU)#(fU)#(mA)#(fC)#(mU)#(mG)#(mA)#(fA) |
| 8154 | MECP2_E1_s_1689 | (mA)#(mA)#(fA)(mA)(fA)(mA)(fA)(mA)(fA)(mA)(mA)(mA)(fA)#(mA)#(mA)-TegChol |
| 8202 | MECP2_E1_as_1689 | P(mU)#(fU)#(mU)(fU)(fU)(fU)(mU)(fU)(mU)(fU)(mU)(fU)(mU)#(fU)#(mU)#(fA)#(mC)#(mU)#(mG)#(fA) |
| 8180 | MECP2_E2_s_113 | (mG)#(mA)#(fG)(mG)(fA)(mG)(fA)(mG)(fA)(mC)(mU)(mG)(fC)#(mU)#(mA)-TegChol |
| 8228 | MECP2_E2_as_113 | P(mU)#(fA)#(mG)(fC)(fA)(fG)(mU)(fC)(mU)(fC)(mU)(fC)(mC)#(fU)#(mC)#(fC)#(mU)#(mC)#(mG)#(fC) |
| 8189 | MECP2_E2_s_128 | (mC)#(mA)#(fU)(mA)(fA)(mA)(fA)(mA)(fU)(mA)(mC)(mA)(fG)#(mA)#(mA)-TegChol |
| 8237 | MECP2_E2_as_128 | P(mU)#(fU)#(mC)(fU)(fG)(fU)(mA)(fU)(mU)(fU)(mU)(fU)(mA)#(fU)#(mG)#(fG)#(mA)#(mG)#(mC)#(fA) |
| 8182 | MECP2_E2_s_131 | (mA)#(mA)#(fA)(mA)(fA)(mU)(fA)(mC)(fA)(mG)(mA)(mC)(fU)#(mC)#(mA)-TegChol |
| 8230 | MECP2_E2_as_131 | P(mU)#(fG)#(mA)(fG)(fU)(fC)(mU)(fG)(mU)(fA)(mU)(fU)(mU)#(fU)#(mU)#(fA)#(mU)#(mG)#(mG)#(fA) |
| 8181 | MECP2_E2_s_146 | (mC)#(mC)#(fA)(mG)(fU)(mU)(fC)(mC)(fU)(mG)(mC)(mU)(fU)#(mU)#(mA)-TegChol |
| 8229 | MECP2_E2_as_146 | P(mU)#(fA)#(mA)(fA)(fG)(fC)(mA)(fG)(mG)(fA)(mA)(fC)(mU)#(fG)#(mG)#(fU)#(mG)#(mA)#(mG)#(fU) |
| 8188 | MECP2_E2_s_147 | (mC)#(mA)#(fG)(mU)(fU)(mC)(fC)(mU)(fG)(mC)(mU)(mU)(fU)#(mG)#(mA)-TegChol |

| | | |
|---|---|---|
| 8236 | MECP2_E2_as_147 | P(mU)#(fC)#(mA)(fA)(fA)(fG)(mC)(fA)(mG)(fG)(mA)(fA)(mC)#(fU)#(mG)#(fG)#(mU)#(mG)#(mA)#(fG) |
| 8184 | MECP2_E2_s_174 | (mC)#(mU)#(fC)(mC)(fC)(mC)(fA)(mG)(fA)(mA)(mU)(mA)(fC)#(mA)#(mA)-TegChol |
| 8232 | MECP2_E2_as_174 | P(mU)#(fU)#(mG)(fU)(fA)(fU)(mU)(fC)(mU)(fG)(mG)(fG)(mG)#(fA)#(mG)#(fU)#(mC)#(mA)#(mC)#(fA) |
| 8179 | MECP2_E2_s_179 | (mC)#(mA)#(fG)(mA)(fA)(mU)(fA)(mC)(fA)(mC)(mC)(mU)(fU)#(mG)#(mA)-TegChol |
| 8227 | MECP2_E2_as_179 | P(mU)#(fC)#(mA)(fA)(fG)(fG)(mU)(fG)(mU)(fA)(mU)(fU)(mC)#(fU)#(mG)#(fG)#(mG)#(mG)#(mA)#(fG) |
| 8178 | MECP2_E2_s_181 | (mG)#(mA)#(fA)(mU)(fA)(mC)(fA)(mC)(fC)(mU)(mU)(mG)(fC)#(mU)#(mA)-TegChol |
| 8226 | MECP2_E2_as_181 | P(mU)#(fA)#(mG)(fC)(fA)(fA)(mG)(fG)(mU)(fG)(mU)(fA)(mU)#(fU)#(mC)#(fU)#(mG)#(mG)#(mG)#(fG) |
| 8183 | MECP2_E2_s_213 | (mC)#(mA)#(fG)(mG)(fA)(mU)(fU)(mC)(fC)(mA)(mU)(mG)(fG)#(mU)#(mA)-TegChol |
| 8231 | MECP2_E2_as_213 | P(mU)#(fA)#(mC)(fC)(fA)(fU)(mG)(fG)(mA)(fA)(mU)(fC)(mC)#(fU)#(mG)#(fU)#(mU)#(mG)#(mG)#(fA) |
| 8186 | MECP2_E2_s_214 | (mA)#(mG)#(fG)(mA)(fU)(mU)(fC)(mC)(fA)(mU)(mG)(mG)(fU)#(mA)#(mA)-TegChol |
| 8234 | MECP2_E2_as_214 | P(mU)#(fU)#(mA)(fC)(fC)(fA)(mU)(fG)(mG)(fA)(mA)(fU)(mC)#(fC)#(mU)#(fG)#(mU)#(mU)#(mG)#(fG) |
| 8185 | MECP2_E2_s_233 | (mG)#(mA)#(fU)(mG)(fU)(mU)(fA)(mG)(fG)(mG)(mC)(mU)(fC)#(mA)#(mA)-TegChol |
| 8233 | MECP2_E2_as_233 | P(mU)#(fU)#(mG)(fA)(fG)(fC)(mC)(fC)(mU)(fA)(mA)(fC)(mA)#(fU)#(mC)#(fC)#(mC)#(mA)#(mG)#(fC) |
| 8187 | MECP2_E2_s_234 | (mA)#(mU)#(fG)(mU)(fU)(mA)(fG)(mG)(fG)(mC)(mU)(mC)(fA)#(mG)#(mA)-TegChol |
| 8235 | MECP2_E2_as_234 | P(mU)#(fC)#(mU)(fG)(fA)(fG)(mC)(fC)(mC)(fU)(mA)(fA)(mC)#(fA)#(mU)#(fC)#(mC)#(mC)#(mA)#(fG) |
| 8201 | MECP2_E2_s_3514 | (mA)#(mA)#(fU)(mG)(fG)(mC)(fA)(mA)(fU)(mG)(mU)(mU)(fU)#(mU)#(mA)-TegChol |
| 8249 | MECP2_E2_as_3514 | P(mU)#(fA)#(mA)(fA)(fA)(fC)(mA)(fU)(mU)(fG)(mC)(fC)(mA)#(fU)#(mU)#(fC)#(mA)#(mA)#(mG)#(fA) |
| 8197 | MECP2_E2_s_3754 | (mU)#(mA)#(fU)(mA)(fU)(mC)(fU)(mA)(fA)(mA)(mU)(mC)(fU)#(mG)#(mA)-TegChol |
| 8245 | MECP2_E2_as_3754 | P(mU)#(fC)#(mA)(fG)(fA)(fU)(mU)(fU)(mA)(fG)(mA)(fU)(mA)#(fU)#(mA)#(fA)#(mG)#(mA)#(mG)#(fA) |

| | | |
|---|---|---|
| 8193 | MECP2_E2_s_4431 | (mU)#(mU)#(fU)(mU)(fU)(mA)(fU)(mG)(fU)(mA)(mU)(mU)(fA)#(mU)#(mA)-TegChol |
| 8241 | MECP2_E2_as_4431 | P(mU)#(fA)#(mU)(fA)(fA)(fU)(mA)(fC)(mA)(fU)(mA)(fA)(mA)#(fA)#(mA)#(fC)#(mC)#(mC)#(mA)#(fA) |
| 8200 | MECP2_E2_s_5875 | (mG)#(mC)#(fA)(mU)(fA)(mU)(fA)(mC)(fA)(mU)(mU)(mU)(fU)#(mU)#(mA)-TegChol |
| 8248 | MECP2_E2_as_5875 | P(mU)#(fA)#(mA)(fA)(fA)(fA)(mU)(fG)(mU)(fA)(mU)(fA)(mU)#(fG)#(mC)#(fC)#(mC)#(mA)#(mA)#(fA) |
| 8190 | MECP2_E2_s_5876 | (mC)#(mA)#(fU)(mA)(fU)(mA)(fC)(mA)(fU)(mU)(mU)(mU)(fU)#(mA)#(mA)-TegChol |
| 8238 | MECP2_E2_as_5876 | P(mU)#(fU)#(mA)(fA)(fA)(fA)(mA)(fU)(mG)(fU)(mA)(fU)(mA)#(fU)#(mG)#(fC)#(mC)#(mC)#(mA)#(fA) |
| 8198 | MECP2_E2_s_5877 | (mA)#(mU)#(fA)(mU)(fA)(mC)(fA)(mU)(fU)(mU)(mU)(mU)(fA)#(mG)#(mA)-TegChol |
| 8246 | MECP2_E2_as_5877 | P(mU)#(fC)#(mU)(fA)(fA)(fA)(mA)(fA)(mU)(fG)(mU)(fA)(mU)#(fA)#(mU)#(fG)#(mC)#(mC)#(mC)#(fA) |
| 8199 | MECP2_E2_s_6278 | (mC)#(mA)#(fG)(mA)(fA)(mA)(fA)(mU)(fU)(mA)(mC)(mA)(fU)#(mU)#(mA)-TegChol |
| 8247 | MECP2_E2_as_6278 | P(mU)#(fA)#(mA)(fU)(fG)(fU)(mA)(fA)(mU)(fU)(mU)(fU)(mC)#(fU)#(mG)#(fC)#(mC)#(mA)#(mA)#(fA) |
| 8196 | MECP2_E2_s_7175 | (mU)#(mA)#(fU)(mU)(fG)(mC)(fA)(mC)(fA)(mA)(mU)(mU)(fA)#(mU)#(mA)-TegChol |
| 8244 | MECP2_E2_as_7175 | P(mU)#(fA)#(mU)(fA)(fA)(fU)(mU)(fG)(mU)(fG)(mC)(fA)(mA)#(fU)#(mA)#(fU)#(mA)#(mC)#(mA)#(fG) |
| 8195 | MECP2_E2_s_7237 | (mA)#(mA)#(fU)(mU)(fA)(mU)(fA)(mC)(fC)(mU)(mG)(mU)(fU)#(mG)#(mA)-TegChol |
| 8243 | MECP2_E2_as_7237 | P(mU)#(fC)#(mA)(fA)(fC)(fA)(mG)(fG)(mU)(fA)(mU)(fA)(mA)#(fU)#(mU)#(fU)#(mU)#(mA)#(mA)#(fC) |
| 8191 | MECP2_E2_s_8813 | (mG)#(mU)#(fG)(mA)(fA)(mA)(fG)(mG)(fA)(mA)(mU)(mU)(fU)#(mU)#(mA)-TegChol |
| 8239 | MECP2_E2_as_8813 | P(mU)#(fA)#(mA)(fA)(fA)(fU)(mU)(fC)(mC)(fU)(mU)(fU)(mC)#(fA)#(mC)#(fC)#(mC)#(mA)#(mC)#(fC) |
| 8194 | MECP2_E2_s_8893 | (mC)#(mA)#(fG)(mA)(fG)(mA)(fC)(mA)(fA)(mA)(mU)(mA)(fU)#(mU)#(mA)-TegChol |
| 8242 | MECP2_E2_as_8893 | P(mU)#(fA)#(mA)(fU)(fA)(fU)(mU)(fU)(mG)(fU)(mC)(fU)(mC)#(fU)#(mG)#(fG)#(mA)#(mA)#(mC)#(fA) |

| | | |
|---|---|---|
| 8192 | MECP2_E2_s_9967 | (mU)#(mU)#(fU)(mG)(fG)(mG)(fA)(mC)(fA)(mA)(mU)(mU)(fA)#(mC)#(mA)-TegChol |
| 8240 | MECP2_E2_as_9967 | P(mU)#(fG)#(mU)(fA)(fA)(fU)(mU)(fG)(mU)(fC)(mC)(fC)(mA)#(fA)#(mA)#(fA)#(mC)#(mA)#(mC)#(fA) |

P=phosphate, #=phosphorothioate, m=2'-Omethyl, f=2'-fluoro, A=adenine, U=uracil, G=guanine, C=cytosine, TegChol = cholesterol conjugate with linker
